# IDR2D identifies reproducible genomic interactions

**DOI:** 10.1101/691295

**Authors:** Konstantin Krismer, Yuchun Guo, David K. Gifford

## Abstract

Chromatin interaction data from protocols such as ChIA-PET, HiChIP, and HiC provide valuable insights into genome organization and gene regulation, but can include spurious interactions that do not reflect underlying genome biology. We introduce a generalization of the Irreproducible Discovery Rate (IDR) method called IDR2D that identifies replicable interactions shared by chromatin interaction experiments. IDR2D provides a principled set of interactions and eliminates artifacts from single experiments. The method is available as a Bioconductor package for the R community, as well as an online service at https://idr2d.mit.edu.

## I. Introduction

The Irreproducible Discovery Rate [1] (IDR) method identifies a robust set of findings that comprise the signal component shared by two replicate experiments. The IDR method is akin to the false discovery rate (FDR) in multiple hypothesis testing, but instead of alleviating alpha error accumulation within one replicate, IDR quantifies the reproducibility of findings using a copula mixture model with one reproducible and one irreproducible component. A finding’s IDR is the probability it belongs to the irreproducible component. This permits findings that are likely in the irreproducible noise component to be eliminated for subsequent analyses. Assessing the IDR of genomic findings has been adopted by ENCODE [2], and is recommended for all ChIP-seq experiments with replicates. IDR has also been used in numerous projects outside of ENCODE [3–6].

Chromatin interaction experiments such as ChIA-PET [7], HiChIP [8], and HiC [9] provide important chromatin structure and gene regulation information, but the complexity of their results and the sampling noise present in their protocols makes the principled analysis of resulting data important. Single replicate false discovery rate (FDR) methods are often used to identify chromatin interactions, but questions can remain about the veracity of the interactions identified as significant as they may not be observed in replicate experiments.

Here we generalize IDR from one dimensional analysis, performed on a single genome coordinate, to the analysis of interactions that are identified in two dimensions by a pair of genome coordinates. We call this extended method Irreproducible Discovery Rate for Two Dimensions (IDR2D) and it can be readily applied to any experimental data type that produces two-dimensional genomic results that admit appropriate distance metrics. We demonstrate the application of IDR2D to data from ChIA-PET, HiChIP, and HiC experiments.

Like IDR, IDR2D independently ranks the findings from each replicate. This ranking reflects the confidence of the finding, with high-confidence interactions at the top and low-confidence interactions at the bottom of the list. In a subsequent step, corresponding interactions between replicates are identified. A genomic interaction from replicate 1 is said to correspond to an interaction in replicate 2, if both their interaction anchors overlap (see figure 1C). After corresponding interactions are identified and ambiguous mappings of interactions between replicates are resolved (see equation 1 and figure 1D), IDRs are computed for each replicated interaction.

**Fig. 1.**
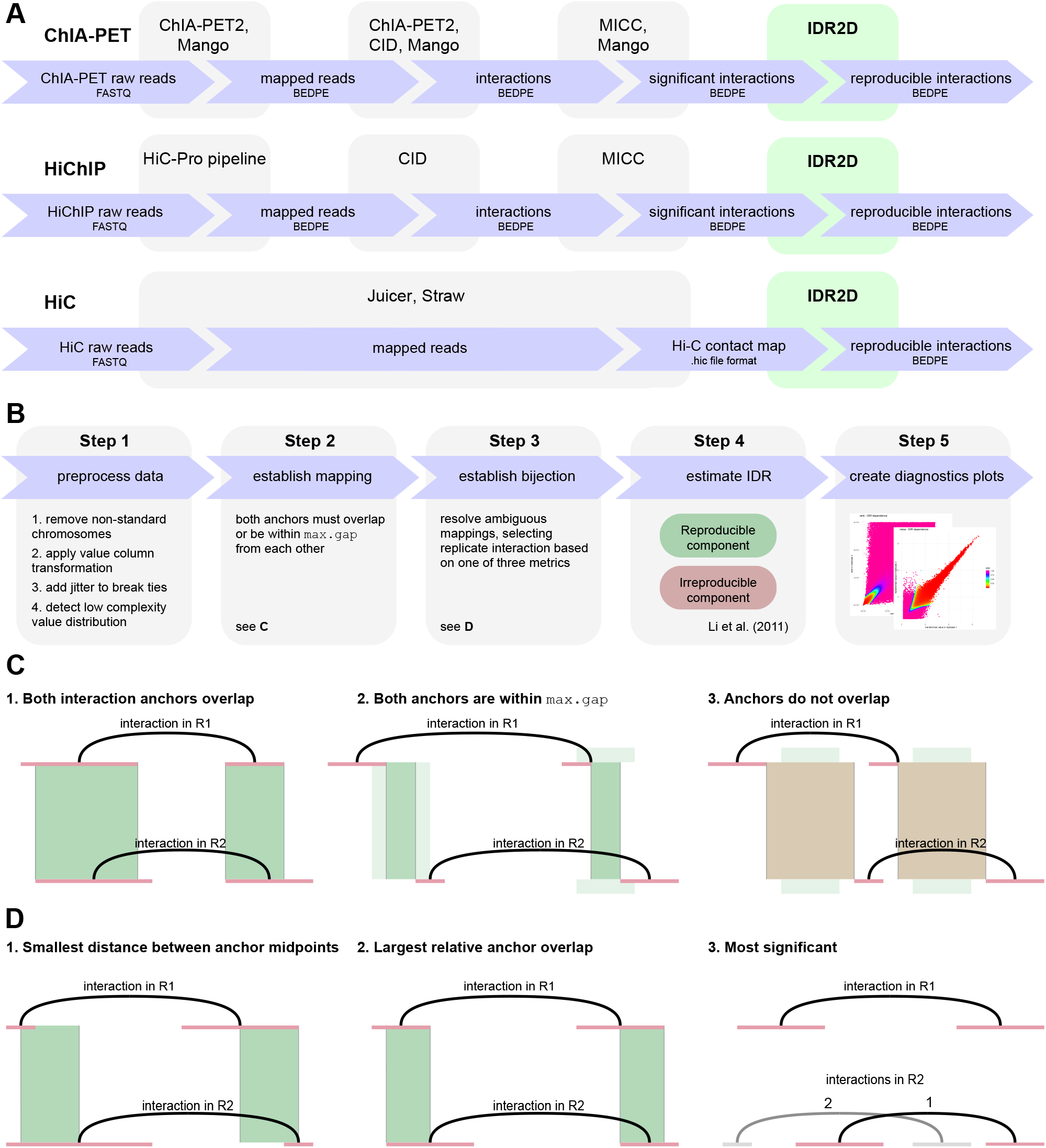
IDR2D identifies reproducible genomic interactions. (**A**) IDR2D is a potential post-processing step in the data analysis pipelines for ChIA-PET, HiChIP, and HiC experiments that were done in replicate. It is compatible with a range of different interaction callers, such as ChIA-PET2, Mango, and CID. (**B**) This schematic depicts the five steps of the IDR2D procedure. In step 1, the data is prepared for IDR analysis, which includes the removal of interactions on non-standard chromosomes and a suitable transformation of the value column, which will be the basis of the ranking. In step 2, interactions in replicate 1 that overlap interactions in replicate 2 are identified, and in step 3 a one-to-one correspondence between overlapping interactions is established by resolving ambiguous cases. After this unambiguous mapping is established, in step 4 the irreproducible discovery rates are estimated for each interaction pair. Lastly, diagnostics plots are created in step 5. (**C**) An interaction in replicate 1 (R1) is assigned to all interactions in R2 for which both interaction anchors overlap or are within `maximum gap` of each other. (**D**) If more than one interaction in R2 overlaps with an interaction in R1, there are three methods to resolve this ambiguous mapping: select the interaction in R2 with (1) the smallest distance between the anchor midpoints (width of the green bars), (2) the largest relative overlap (width of the green bars divided by the sum of the anchor widths), or (3) the lowest rank sum of the interactions, which prioritizes more significant interactions.

If interaction *x*_1_ in replicate 1 overlaps with more than one interaction in replicate 2, the ambiguous mapping is resolved by choosing 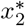 in the following way:

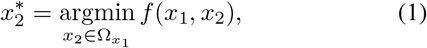

where 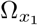 is the set of interactions in replicate 2 that overlaps with the interaction *x*_1_ in replicate 1, and *f* (⋅, ⋅) is the *ambiguity resolution value* (ARV) between an interaction in replicate 1 and an overlapping interaction in replicate 2. Depending on the ambiguity resolution method, this value corresponds to the genomic distance between anchor midpoints (see 1. in figure 1D), the additive inverse of the relative anchor overlap (see 2. in figure 1D), or the sum of the interaction ranks, where more significant interactions have lower ranks.

IDR2D is used as the final step in chromatin interaction data workflows (see figure 1A). The input to IDR2D are BEDPE formatted files of genomic interactions, where each genomic interaction has a score associated with it. This score is used to rank the interactions and can be probability-based, such as the scores from MICC-based methods [10–12], or based on a heuristic. Figure 1B breaks the IDR2D procedure into five steps.

## II. Materials and Methods

### A. IDR

IDR2D extends the reference implementation of IDR in R by Qunhua Li [1]. All datasets were analyzed with the IDR2D package using default parameters. We used *overlap* as ambiguity resolution method and allowed no gaps between overlapping interactions (`max.gap = 0L`). The applied value transformations were dependent on the interaction calling method. The results were not sensitive to the initial values of the mean, standard deviation, correlation coefficient, or proportion of the reproducible component.

### B. ChIA-PET datasets

We used 17 ChIA-PET datasets associated with protein factors that include POL2RA, CTCF and RAD21 from selected cell types (Supplementary Table S1). The datasets were downloaded from the ENCODE Project portal (https://www.encodeproject.org/). All FASTQ files of both biological replicates were pre-processed and aligned to the hg19 genome assembly using steps 1, 2 and 3 in the ChIA-PET data analysis software *Mango*.

### C. HiChIP datasets

The FASTQ SMC1A HiChIP data from GM12878 cells [8] were downloaded from the NCBI GEO portal (GSE80820). Raw read files were analyzed with HiC-Pro [13], and interactions were subsequently called by CID and *hichipper* [14].

### D. HiC datasets

Preprocessed contact matrix files in *.hic* format were downloaded from the NCBI GEO portal (see supplementary table S1 for details) and parsed with *Straw*, a data extraction API for *.hic* files [15].

### E. Mango pipeline

Mango 1.2.1 [16] was downloaded from https://github.com/dphansti/mango. Additionally, we installed the dependencies *R* 3.4.4, *bedtools* 2.26.0, *macs2* (version 2.1.1.20160309) [17], and *bowtie-align* 1.2 [18]. Mango was executed with the default parameters and the flags `verboseoutput` and `reportallpairs` were set. For datasets that were generated with the ChIA-PET Tn5 tagmentation protocol, additional parameters recommended by the author were used: `-keepempty TRUE -maxlength 1000 -shortreads FALSE`. For subsequent IDR2D analyses, we used the P column in the Mango output files to establish the ranking of interactions. This column contains unadjusted p-values, which were transformed using the `log.additive.inverse` transformation to match the IDR semantics of the value column, where interactions with larger values are more likely to be true interactions.

The BEDPE files generated by Mango after step 3 were also used by the ChIA-PET2 and CID pipelines.

### F. ChIA-PET2 pipeline

*ChIA-PET2* 0.9.2 [11] was obtained from https://github.com/GuipengLi/ChIA-PET2. The default setting was used for all parameters, except that the starting step was set to 4 to start the analysis from Mango-derived BEDPE files. The ranking for the IDR2D analysis was established by the untransformed `-log10(1-PostProb)` column, which is an output of *MICC* [10], a Bayesian mixture model used internally by ChIA-PET2 and *CID*.

### G. CID pipeline

CID 1.0 [12] is part of the Java genomics software package *GEM* 3.4, which was downloaded from http://groups.csail.mit.edu/cgs/gem/versions.html. We used the default CID parameters. Before running MICC, we filtered all interactions that were supported by only one PET read. Same as with ChIA-PET2, we used the untransformed `-log10(1-PostProb)` column to rank interactions in IDR2D.

### H. Package and web development

The R package development process was supported using *devtools*. We used *roxygen2* for inline function documentation, and *knitr* and *R Markdown* for package vignettes. With the R package we provide a platform-independent implementation of the methods introduced in this paper. The HiC analysis part of the package requires the Python package *hic-straw* [15], which is a data extraction API for HiC contact maps.

The website was developed in R with the reactive web application framework *shiny* from RStudio. The components of the graphical user interface were provided by shiny and *shinyBS*, which serve as an R wrapper for the components of the Bootstrap front-end web development framework. The analysis job queue of the website uses an *SQLite* database.

## III. Results

### A. IDR2D identifies reproducible components of ENCODE ChIA-PET experiments

To assess the performance and utility of IDR2D we analyzed the read data of 17 ChIA-PET experiments that had replicates (see Supplementary Table S1 for details). Mango was used for data preprocessing such as linker removal, read mapping (via bowtie), and peak calling (via macs2). We called interactions with three different methods (ChIA-PET2, CID, and Mango) and then used IDR2D to identify reproducible interactions across replicates. The number of identified interactions varies greatly between the three interaction callers, with on average CID identifying the most, and Mango the fewest interactions (see figure 2C). The overall reproducibility of interactions is strongly dependent on the experiment and less dependent on the interaction caller used. For example, the ChIA-PET experiments `Snyder.GM19239.RAD21` and `Snyder.GM19240.RAD21` show poor reproducibility across all three interaction calling methods. By identifying the reproducible component within each of the replicated experiments, IDR2D helps to assess the overall reproducibility of each experiment, as well as the reproducibility of individual findings, which in turn informs the conclusions drawn from the data. In addition, it can be used to help qualify new experimental protocols for consistent results.

**Fig. 2.**
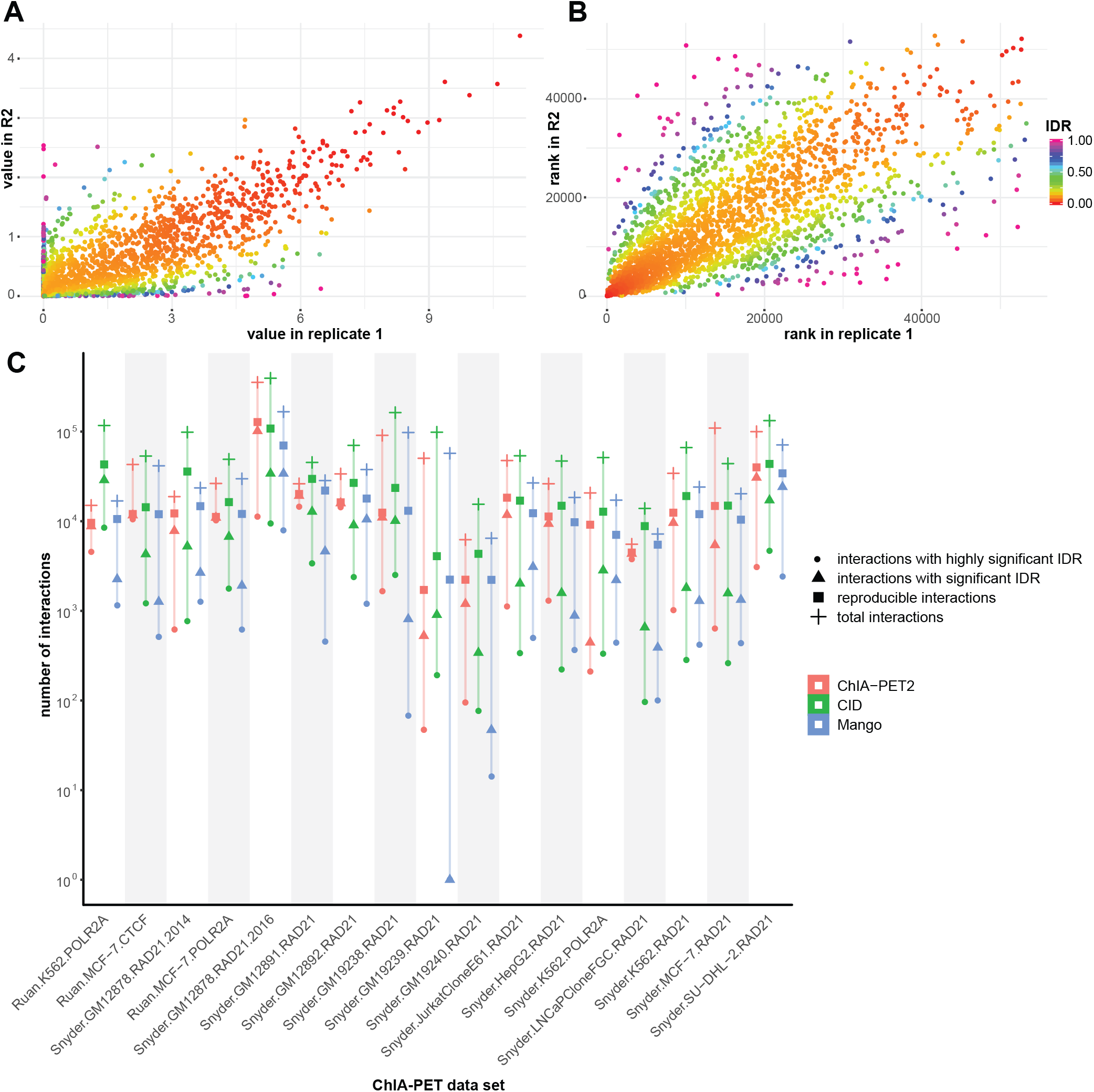
IDR2D analysis of 17 replicated ChIA-PET experiments identifies reproducible components. (**A**) Diagnostic scatterplot of IDRs of genomic interactions called by CID from replicated ChIA-PET experiments targeting RAD21 in HepG2 cells. Plotted are replicated interactions with their estimated IDR (color) and their scores in the two replicates (position). As expected, interactions with low IDRs that have a low probability of belonging to the irreproducible component, are along the diagonal (similar scores in both replicates). (**B**) Similar to panel **A**, but plots interaction ranks instead of scores (higher score results in lower rank). (**C**) A comparison of ChIA-PET interaction callers ChIA-PET2, CID, and Mango across 17 ChIA-PET experiments. Significant IDR *<* 0.05, highly significant IDR *<* 0.01, *total interactions* is the number of interactions in replicate 1.

Furthermore, we used IDR2D to analyze experimental results from replicated HiChIP (see supplemental tables S5 and S6). Similar to ChIA-PET, IDR2D can identify reproducible HiChIP interactions and expose poorly replicated experiments, which is valuable information for subsequent analysis steps.

### B. Mappings of genomic interactions between replicated ChIA-PET experiments are predominantly unambiguous

The great majority of interactions in replicate 1 that overlapped with interactions in replicate 2 overlapped with only one interaction, leading to an unambiguous assignment of corresponding replicated interactions (see figure 3). Unsurprisingly, the number of ambiguous mappings (interactions in replicate 1 that overlap with more than one interaction in replicate 2) increases when the *maximum gap* is increased, the tolerated distance between anchors that are considered to overlap. On average, only 2.66% of interactions are ambiguous in the case of zero *max gap*, whereas this number increases to 8.00% and 24.11% with maximum gaps of 1000 and 5000 bp, respectively.

**Fig. 3.**
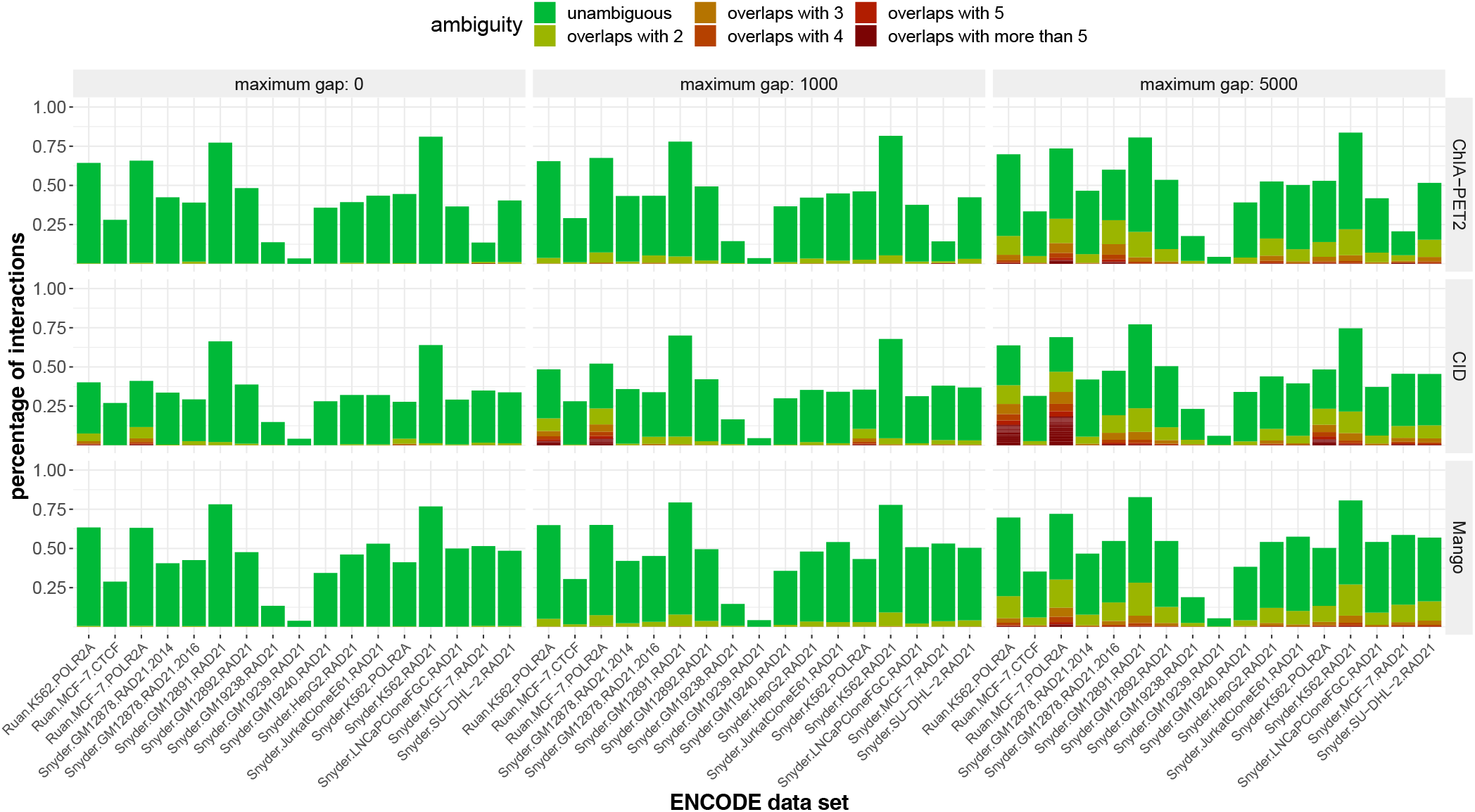
Mappings of genomic interactions between replicated ChIA-PET experiments are predominantly unambiguous. The great majority of interactions in replicate 1 that overlapped with interactions in replicate 2 only overlapped with one interaction, leading to an unambiguous assignment of corresponding replicated interactions (green bars). Unsurprisingly, the number of ambiguous mappings (interactions in replicate 1 that overlap with more than one interaction in replicate 2) increases when the maximum acceptable gap is increased, the tolerated distance between anchors to still be considered overlapping.

There are more ambiguous mappings between replicated interactions that were called with CID (14.73% for CID, 9.90% for ChIA-PET2, and 10.14% for Mango). We expect this is because (1) CID on average calls significantly more interactions than ChIA-PET2 and Mango, and (2) interactions called with CID exhibit a wider range of anchor lengths, and longer anchors naturally increase the probability of overlap.

### C. Assessing reproducibility of HiC experiments

When analyzing pairs of HiC experiments with IDR2D, blocks from HiC contact matrices are used as observations. The resolution of contact matrix values typically ranges between 5 kbp (kilo base pairs) to 2.5 Mbp blocks. With the fixed grid of contact map observations, finding corresponding observations in the second replicate is straightforward. Each block in replicate 1 is simply matched with the block spanning the same genomic regions in replicate 2. Blocks are subsequently ranked by their read counts and analyzed using the same procedure that was used for ChIA-PET and HiChIP data.

In figure 4 we show IDR2D results for three pairs of HiC experiments. The first pair of HiC experiments consists of true replicate experiments in GM12878 cells [19]. The second pair of experiments were obtained in phased murine embryonic kidney fibroblasts, where allele-specific HiC reads [19] were available (*different alleles* in figure 4) [20], and the third pair of HiC experiments were obtained before and after the overexpression of NSD2 (different treatments) [21]. GEO identifiers of all data sets are listed in the supplemental table S1 and detailed results in supplemental table S7. Figure 4A gives an overview of all data sets and all resolutions, showing that, as expected, the reproducibility is highest between true replicates, and in general higher at lower resolutions (larger blocks). Figure 4B depicts the distribution of IDR values for chromosome 1 of each of the HiC pairs at block resolutions of 1 Mbp, 250 kbp, and 10 kbp. The largest fraction of reproducible blocks is found between replicated experiments. Figures 4C-E are scatterplots of interaction pairs (corresponding blocks in the contact matrices) of the two experiments, where the color denotes the IDR value of the interaction pair.

**Fig. 4.**
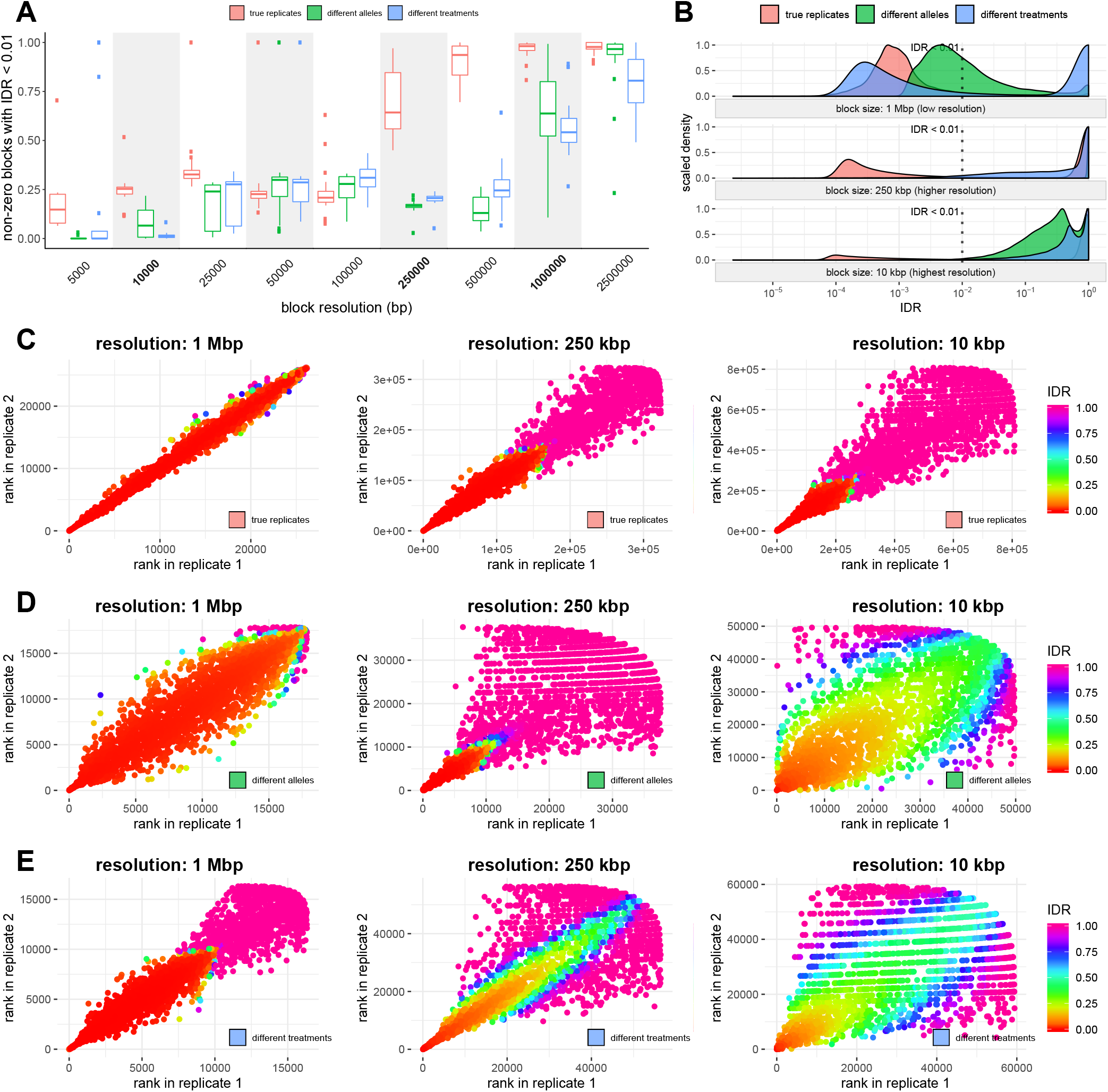
Reproducibility analysis of HiC experiments. (**A**) Summary of IDR2D results on individual chromosomes of three pairs of HiC experiments, True replicate HiC experiments (Lieberman.GM12878) are compared to IDR2D analysis of HiC experiments of different alleles (Lieberman.Patski) and different treatments (Skok.NSD2). (**B**) Histograms of the IDR distribution of IDR values for all blocks of chromosome 1 for the three pairs of HiC experiments. (**C**) Scatterplots of block ranks of chromosome 1 of the two HiC replicate experiments, colored by IDR. (**D**) Analagous to **C**, for HiC experiments of paternal and maternal alleles. (**E**) Analagous to **C**, for HiC experiments before and after overexpression of NSD2. Axis scales are not fixed between scatterplots.

In addition to computing IDR values, IDR2D produces diagnostic plots that interpret the overall reproducibility of a pair of HiC experiments, as well as identify reproducible parts of HiC contact matrices for a more focused, downstream analysis.

## IV. Discussion

IDR2D offers a complementary way to evaluate the results of chromatin interaction experiments for significance, and provides a foundation for subsequent analysis such as enhancer-gene mapping that incorporates the important concept of experimental replicability.

## Availability

The implementation of IDR2D facilitates workflow integration with other data analysis pipelines, and is also web-accessible at https://idr2d.mit.edu. IDR2D is implemented in R and bundled as an R/Bioconductor package (idr2d), supporting observations with both one-dimensional and two-dimensional genomic coordinates. The IDR2D website implementation offers a number of ways to transform the scores to match IDR requirements, and to map interactions between replicates. The source code of the R package is hosted on GitHub (https://github.com/gifford-lab/idr2d).

## Author contributions

Conceptualization, D.K.G.; Methodology, K.K.; Software, K.K.; Formal Analysis, K.K., and Y.G.; Investigation, K.K., and Y.G.; Resources, D.K.G.; Data Curation, K.K., and Y.G.; Writing - Original Draft, K.K.; Writing - Review & Editing, K.K., Y.G., and D.K.G.; Visualization, K.K.; Supervision, D.K.G.; Funding Acquisition, D.K.G.

## Acknowledgements

We thank members of the Gifford lab for insightful suggestions and discussions. We also thank William Stafford Noble for his thoughtful comments on an earlier version of the manuscript.

