## Supplemental Material for "IDR2D identifies reproducible genomic interactions"

| ChIA-PET dataset identifier | Target | Cell line | ENCODE identifier |
|---|---|---|---|
| Ruan.K562.POLR2A | POLR2A | K562 | ENCSR000BZY |
| Ruan.MCF-7.POLR2A | POLR2A | MCF-7 | ENCSR000CAA |
| Snyder.K562.POLR2A | POLR2A | K562 | ENCSR000FDC |
| Ruan.MCF-7.CTCF | CTCF | MCF-7 | ENCSR000CAD |
| Snyder.GM12878.RAD21.2014 | RAD21 | GM12878 | ENCSR752QCX |
| Snyder.GM12878.RAD21.2016 | RAD21 | GM12878 | ENCSR981FNA |
| Snyder.GM12891.RAD21 | RAD21 | GM12891 | ENCSR299VMZ |
| Snyder.GM12892.RAD21 | RAD21 | GM12892 | ENCSR033GUP |
| Snyder.GM19238.RAD21 | RAD21 | GM19238 | ENCSR527RXH |
| Snyder.GM19239.RAD21 | RAD21 | GM19239 | ENCSR479MTN |
| Snyder.GM19240.RAD21 | RAD21 | GM19240 | ENCSR312TUD |
| Snyder.HepG2.RAD21 | RAD21 | HepG2 | ENCSR014ZXR |
| Snyder.JurkatCloneE61.RAD21 | RAD21 | Jurkat clone E61 | ENCSR465NNU |
| Snyder.K562.RAD21 | RAD21 | K562 | ENCSR000FDB |
| Snyder.LNCaPCloneFGC.RAD21 | RAD21 | LNCaP clone FGC | ENCSR011ITK |
| Snyder.MCF-7.RAD21 | RAD21 | MCF-7 | ENCSR716WZI |
| Snyder.SU-DHL-2.RAD21 | RAD21 | SU-DHL-2 | ENCSR466AXT |
| **HiChIP dataset identifier** | **Target** | **Cell line** | **GEO identifier** |
| Chang.GM12878.H3K27ac | H3K27ac | GM12878 | GSE101498 |
| Chang.K562.H3K27ac | H3K27ac | K562 | GSE101498 |
| Chang.mES.H3K27ac | H3K27ac | mES | GSE101498 |
| Chang.MyLa.H3K27ac | H3K27ac | MyLa | GSE101498 |
| Chang.Naive.CTCF | CTCF | Naive | GSE101498 |
| Chang.Naive.H3K27ac | H3K27ac | Naive | GSE101498 |
| Chang.Th17.H3K27ac | H3K27ac | Th17 | GSE101498 |
| Chang.Treg.H3K27ac | H3K27ac | Treg | GSE101498 |
| Flynn.GM12878.Smc1a | Smc1a | GM12878 | GSE80820 |
| **HiC dataset identifier** | | **Cell line** | **GEO identifier** |
| Lieberman.GM12878 | | GM12878 | GSE63525 |
| Lieberman.Patski | | Patski | GSE71831 |
| Skok.NSD2 | | multiple myeloma | GSE131651 |

Table S1: ChIA-PET, HiChIP, and HiC datasets used in this study.

| Dataset identifier | Total int. | Rep. int. | IDR < 0.05 | IDR < 0.01 |
|---|---|---|---|---|
| Ruan.K562.POLR2A | 15,029 | 9594 | 8816 | 4919 |
| Ruan.MCF-7.POLR2A | 23,540 | 14,718 | 2714 | 1251 |
| Snyder.K562.POLR2A | 20,683 | 9142 | 446 | 220 |
| Ruan.MCF-7.CTCF | 42,958 | 12,042 | 11,664 | 10,503 |
| Snyder.GM12878.RAD21.2014 | 26,376 | 11,176 | 10,977 | 10,191 |
| Snyder.GM12878.RAD21.2016 | 354,536 | 127,172 | 101,330 | 11,164 |
| Snyder.GM12891.RAD21 | 26,180 | 20,144 | 19,245 | 14,628 |
| Snyder.GM12892.RAD21 | 33,731 | 16,253 | 15,906 | 14,401 |
| Snyder.GM19238.RAD21 | 90,887 | 12,471 | 11,031 | 1676 |
| Snyder.GM19239.RAD21 | 50,387 | 1712 | 526 | 47 |
| Snyder.GM19240.RAD21 | 6225 | 2226 | 1212 | 94 |
| Snyder.HepG2.RAD21 | 47,544 | 18,308 | 12,819 | 1300 |
| Snyder.JurkatCloneE61.RAD21 | 18,408 | 9745 | 889 | 367 |
| Snyder.K562.RAD21 | 5540 | 4470 | 4352 | 3782 |
| Snyder.LNCaPCloneFGC.RAD21 | 34,260 | 12,456 | 9718 | 1056 |
| Snyder.MCF-7.RAD21 | 109,485 | 14,793 | 5117 | 639 |
| Snyder.SU-DHL-2.RAD21 | 99,864 | 39,891 | 31,437 | 3266 |

Table S2: IDR2D analysis of ChIA-PET interactions called by ChIA-PET2. Columns are (1) *total number of interactions in replicate 1*, (2) *number of reproducible interactions*, (3) *number of reproducible interactions with IDR* $< 0.05$, and (4) *number of reproducible interactions with IDR* $< 0.01$.

| Dataset identifier | Total int. | Rep. int. | IDR < 0.05 | IDR < 0.01 |
|---|---|---|---|---|
| Ruan.K562.POLR2A | 116,932 | 40,693 | 26,958 | 8698 |
| Ruan.MCF-7.POLR2A | 98,484 | 33,147 | 5064 | 681 |
| Snyder.K562.POLR2A | 51,323 | 12,352 | 6300 | 1513 |
| Ruan.MCF-7.CTCF | 53,351 | 14,321 | 4163 | 1162 |
| Snyder.GM12878.RAD21.2014 | 49,102 | 16,309 | 6556 | 1733 |
| Snyder.GM12878.RAD21.2016 | 394,045 | 105,704 | 36,662 | 10,363 |
| Snyder.GM12891.RAD21 | 45,295 | 29,509 | 13,398 | 3633 |
| Snyder.GM12892.RAD21 | 70,222 | 26,637 | 9594 | 2457 |
| Snyder.GM19238.RAD21 | 162,936 | 23,444 | 10,573 | 2745 |
| Snyder.GM19239.RAD21 | 98,449 | 4074 | 169 | 22 |
| Snyder.GM19240.RAD21 | 15,473 | 4325 | 295 | 65 |
| Snyder.HepG2.RAD21 | 53,725 | 16,886 | 1876 | 267 |
| Snyder.JurkatCloneE61.RAD21 | 46,965 | 14,837 | 1642 | 228 |
| Snyder.K562.RAD21 | 13,891 | 8791 | 790 | 104 |
| Snyder.LNCaPCloneFGC.RAD21 | 66,356 | 19,021 | 2013 | 299 |
| Snyder.MCF-7.RAD21 | 43,930 | 14,754 | 1641 | 262 |
| Snyder.SU-DHL-2.RAD21 | 132,610 | 43,197 | 17,460 | 4765 |

Table S3: IDR2D analysis of ChIA-PET interactions called by CID. Columns are (1) *total number of interactions in replicate 1*, (2) *number of reproducible interactions*, (3) *number of reproducible interactions with IDR* $< 0.05$, and (4) *number of reproducible interactions with IDR* $< 0.01$.

| Dataset identifier | Total int. | Rep. int. | IDR < 0.05 | IDR < 0.01 |
|---|---|---|---|---|
| Ruan.K562.POLR2A | 16,847 | 10,581 | 2273 | 1149 |
| Ruan.MCF-7.POLR2A | 23,540 | 14,718 | 2714 | 1251 |
| Snyder.K562.POLR2A | 17,191 | 7051 | 2419 | 477 |
| Ruan.MCF-7.CTCF | 41,542 | 11,975 | 1274 | 502 |
| Snyder.GM12878.RAD21.2014 | 29,770 | 12,057 | 1993 | 619 |
| Snyder.GM12878.RAD21.2016 | 166,543 | 69,897 | 36,597 | 8039 |
| Snyder.GM12891.RAD21 | 28,426 | 21,983 | 3976 | 338 |
| Snyder.GM12892.RAD21 | 37,730 | 17,900 | 11,050 | 1182 |
| Snyder.GM19238.RAD21 | 97,679 | 13,084 | 1038 | 109 |
| Snyder.GM19239.RAD21 | 57,254 | 2231 | 1 | 0 |
| Snyder.GM19240.RAD21 | 6462 | 2219 | 47 | 14 |
| Snyder.HepG2.RAD21 | 26,747 | 12,285 | 3091 | 494 |
| Snyder.JurkatCloneE61.RAD21 | 18,408 | 9745 | 889 | 367 |
| Snyder.K562.RAD21 | 7203 | 5474 | 412 | 98 |
| Snyder.LNCaPCloneFGC.RAD21 | 24,065 | 11,992 | 1308 | 420 |
| Snyder.MCF-7.RAD21 | 20,277 | 10,412 | 1453 | 511 |
| Snyder.SU-DHL-2.RAD21 | 71,043 | 34,226 | 24,909 | 2552 |

Table S4: IDR2D analysis of ChIA-PET interactions called by Mango. Columns are (1) *total number of interactions in replicate 1*, (2) *number of reproducible interactions*, (3) *number of reproducible interactions with IDR* $< 0.05$, and (4) *number of reproducible interactions with IDR* $< 0.01$.

| Dataset identifier | Total int. | Rep. int. | IDR < 0.05 | IDR < 0.01 |
|---|---|---|---|---|
| Chang.GM12878.H3K27ac | 8,170,291 | 4,076,304 | 2,133,248 | 408,385 |
| Chang.K562.H3K27ac | 6,059,672 | 2,724,718 | 991,394 | 295,193 |
| Chang.mES.H3K27ac | 4,997,523 | 2,593,482 | 1,902,221 | 118,607 |
| Chang.MyLa.H3K27ac | 6,010,073 | 2,745,081 | 1,052,481 | 345,503 |
| Chang.Naive.CTCF | 1,368,563 | 52,845 | 6885 | 1607 |
| Chang.Naive.H3K27ac | 69,861 | 49,823 | 13,504 | 1821 |
| Chang.Th17.H3K27ac | 2,135,349 | 1,418,902 | 889,736 | 188,608 |
| Chang.Treg.H3K27ac | 365,827 | 253,529 | 105,227 | 15,856 |
| Flynn.GM12878.Smc1a | 6,036,994 | 2,232,884 | 369,344 | 4166 |

Table S5: IDR2D analysis of HiChIP interactions called by CID. Columns are (1) *total number of interactions in replicate 1*, (2) *number of reproducible interactions*, (3) *number of reproducible interactions with IDR* $< 0.05$, and (4) *number of reproducible interactions with IDR* $< 0.01$.

| Dataset identifier | Total int. | Rep. int. | IDR < 0.05 | IDR < 0.01 |
|---|---|---|---|---|
| Flynn.GM12878.Smc1a | 2,312,685 | 1,126,785 | 71,850 | 2892 |

Table S6: IDR2D analysis of HiChIP interactions called by hichipper. Columns are (1) *total number of interactions in replicate 1*, (2) *number of reproducible interactions*, (3) *number of reproducible interactions with IDR* $< 0.05$, and (4) *number of reproducible interactions with IDR* $< 0.01$.

| Dataset identifier | Chr. | Resolution | Non-empty blocks | IDR < 0.05 | IDR < 0.01 |
|---|---|---:|---:|---:|---:|
| Lieberman.GM12878 | chr1 | 5,000 | 810,663 | 195,113 | 180,896 |
| Lieberman.GM12878 | chr1 | 10,000 | 808,144 | 222,766 | 210,847 |
| Lieberman.GM12878 | chr1 | 25,000 | 531,624 | 179,436 | 173,794 |
| Lieberman.GM12878 | chr1 | 50,000 | 491,974 | 112,121 | 110,096 |
| Lieberman.GM12878 | chr1 | 100,000 | 629,037 | 141,128 | 130,959 |
| Lieberman.GM12878 | chr1 | 250,000 | 323,444 | 151,273 | 145,581 |
| Lieberman.GM12878 | chr1 | 500,000 | 100,553 | 84,852 | 76,715 |
| Lieberman.GM12878 | chr1 | 1,000,000 | 26,188 | 25,470 | 24,582 |
| Lieberman.GM12878 | chr1 | 2,500,000 | 4,278 | 4,058 | 3,893 |
| Lieberman.GM12878 | chr2 | 5,000 | 723,520 | 196,615 | 56,908 |
| Lieberman.GM12878 | chr2 | 10,000 | 782,056 | 272,553 | 92,570 |
| Lieberman.GM12878 | chr2 | 25,000 | 554,758 | 182,181 | 176,264 |
| Lieberman.GM12878 | chr2 | 50,000 | 497,542 | 193,676 | 112,630 |
| Lieberman.GM12878 | chr2 | 100,000 | 679,578 | 155,472 | 141,249 |
| Lieberman.GM12878 | chr2 | 250,000 | 373,283 | 230,169 | 188,254 |
| Lieberman.GM12878 | chr2 | 500,000 | 113,927 | 89,447 | 79,260 |
| Lieberman.GM12878 | chr2 | 1,000,000 | 29,243 | 28,656 | 27,907 |
| Lieberman.GM12878 | chr2 | 2,500,000 | 4,753 | 4,672 | 4,591 |
| Lieberman.GM12878 | chr3 | 5,000 | 599,232 | 135,122 | 127,995 |
| Lieberman.GM12878 | chr3 | 10,000 | 640,635 | 168,021 | 159,080 |
| Lieberman.GM12878 | chr3 | 25,000 | 461,102 | 232,605 | 156,520 |
| Lieberman.GM12878 | chr3 | 50,000 | 403,388 | 168,187 | 113,571 |
| Lieberman.GM12878 | chr3 | 100,000 | 549,425 | 139,110 | 41,564 |
| Lieberman.GM12878 | chr3 | 250,000 | 277,284 | 151,653 | 143,743 |
| Lieberman.GM12878 | chr3 | 500,000 | 76,208 | 67,754 | 59,398 |
| Lieberman.GM12878 | chr3 | 1,000,000 | 19,306 | 19,074 | 18,766 |
| Lieberman.GM12878 | chr3 | 2,500,000 | 3,240 | 3,233 | 3,225 |
| Lieberman.GM12878 | chr4 | 5,000 | 420,874 | 121,508 | 34,313 |
| Lieberman.GM12878 | chr4 | 10,000 | 485,037 | 118,935 | 112,651 |
| Lieberman.GM12878 | chr4 | 25,000 | 390,484 | 187,368 | 105,089 |
| Lieberman.GM12878 | chr4 | 50,000 | 325,098 | 89,002 | 87,087 |
| Lieberman.GM12878 | chr4 | 100,000 | 469,066 | 89,047 | 84,236 |
| Lieberman.GM12878 | chr4 | 250,000 | 219,566 | 131,924 | 125,304 |
| Lieberman.GM12878 | chr4 | 500,000 | 70,476 | 51,295 | 49,639 |
| Lieberman.GM12878 | chr4 | 1,000,000 | 17,960 | 15,055 | 14,515 |
| Lieberman.GM12878 | chr4 | 2,500,000 | 2,926 | 2,702 | 2,611 |
| Lieberman.GM12878 | chr5 | 5,000 | 493,042 | 137,013 | 39,869 |
| Lieberman.GM12878 | chr5 | 10,000 | 543,723 | 190,010 | 65,579 |
| Lieberman.GM12878 | chr5 | 25,000 | 405,801 | 204,970 | 134,614 |
| Lieberman.GM12878 | chr5 | 50,000 | 369,247 | 87,537 | 85,909 |
| Lieberman.GM12878 | chr5 | 100,000 | 471,544 | 104,213 | 97,556 |
| Lieberman.GM12878 | chr5 | 250,000 | 206,777 | 118,590 | 113,362 |
| Lieberman.GM12878 | chr5 | 500,000 | 62,465 | 46,734 | 44,794 |

*Continued on next page*

Table S7 – *Continued from previous page*

| Dataset identifier | Chr. | Resolution | Non-empty blocks | IDR < 0.05 | IDR < 0.01 |
|---|---|---|---|---|---|
| Lieberman.GM12878 | chr5 | 1,000,000 | 15,934 | 14,957 | 14,030 |
| Lieberman.GM12878 | chr5 | 2,500,000 | 2,701 | 2,661 | 2,622 |
| Lieberman.GM12878 | chr6 | 5,000 | 533,326 | 144,106 | 41,979 |
| Lieberman.GM12878 | chr6 | 10,000 | 559,146 | 148,886 | 141,479 |
| Lieberman.GM12878 | chr6 | 25,000 | 391,304 | 203,798 | 146,938 |
| Lieberman.GM12878 | chr6 | 50,000 | 335,017 | 85,056 | 83,452 |
| Lieberman.GM12878 | chr6 | 100,000 | 450,897 | 90,657 | 85,894 |
| Lieberman.GM12878 | chr6 | 250,000 | 205,011 | 115,894 | 109,533 |
| Lieberman.GM12878 | chr6 | 500,000 | 56,749 | 53,852 | 50,613 |
| Lieberman.GM12878 | chr6 | 1,000,000 | 14,345 | 14,267 | 14,159 |
| Lieberman.GM12878 | chr6 | 2,500,000 | 2,415 | 2,401 | 2,384 |
| Lieberman.GM12878 | chr7 | 5,000 | 437,592 | 117,112 | 32,835 |
| Lieberman.GM12878 | chr7 | 10,000 | 472,781 | 162,683 | 54,365 |
| Lieberman.GM12878 | chr7 | 25,000 | 346,205 | 171,321 | 112,967 |
| Lieberman.GM12878 | chr7 | 50,000 | 309,624 | 122,117 | 67,032 |
| Lieberman.GM12878 | chr7 | 100,000 | 398,266 | 89,716 | 82,787 |
| Lieberman.GM12878 | chr7 | 250,000 | 172,975 | 90,090 | 86,290 |
| Lieberman.GM12878 | chr7 | 500,000 | 48,503 | 45,505 | 41,819 |
| Lieberman.GM12878 | chr7 | 1,000,000 | 12,445 | 12,341 | 12,186 |
| Lieberman.GM12878 | chr7 | 2,500,000 | 2,080 | 2,043 | 2,013 |
| Lieberman.GM12878 | chr8 | 5,000 | 403,695 | 110,526 | 31,924 |
| Lieberman.GM12878 | chr8 | 10,000 | 443,115 | 113,894 | 107,971 |
| Lieberman.GM12878 | chr8 | 25,000 | 335,194 | 111,901 | 107,842 |
| Lieberman.GM12878 | chr8 | 50,000 | 313,846 | 72,524 | 71,048 |
| Lieberman.GM12878 | chr8 | 100,000 | 392,026 | 91,831 | 85,530 |
| Lieberman.GM12878 | chr8 | 250,000 | 149,140 | 99,873 | 93,884 |
| Lieberman.GM12878 | chr8 | 500,000 | 40,951 | 38,507 | 36,229 |
| Lieberman.GM12878 | chr8 | 1,000,000 | 10,510 | 10,439 | 10,314 |
| Lieberman.GM12878 | chr8 | 2,500,000 | 1,770 | 1,763 | 1,755 |
| Lieberman.GM12878 | chr9 | 5,000 | 360,162 | 93,962 | 25,567 |
| Lieberman.GM12878 | chr9 | 10,000 | 372,945 | 100,735 | 94,393 |
| Lieberman.GM12878 | chr9 | 25,000 | 268,070 | 84,515 | 81,627 |
| Lieberman.GM12878 | chr9 | 50,000 | 230,751 | 92,595 | 45,628 |
| Lieberman.GM12878 | chr9 | 100,000 | 251,298 | 58,406 | 55,179 |
| Lieberman.GM12878 | chr9 | 250,000 | 88,293 | 52,796 | 50,496 |
| Lieberman.GM12878 | chr9 | 500,000 | 25,407 | 23,488 | 22,560 |
| Lieberman.GM12878 | chr9 | 1,000,000 | 6,969 | 6,609 | 6,493 |
| Lieberman.GM12878 | chr9 | 2,500,000 | 1,265 | 1,227 | 1,212 |
| Lieberman.GM12878 | chr10 | 5,000 | 427,941 | 115,314 | 33,459 |
| Lieberman.GM12878 | chr10 | 10,000 | 455,285 | 160,620 | 54,640 |
| Lieberman.GM12878 | chr10 | 25,000 | 322,398 | 108,634 | 104,978 |
| Lieberman.GM12878 | chr10 | 50,000 | 309,327 | 105,336 | 47,690 |

*Continued on next page*

Table S7 – *Continued from previous page*

| Dataset identifier | Chr. | Resolution | Non-empty blocks | IDR < 0.05 | IDR < 0.01 |
|---|---|---|---|---|---|
| Lieberman.GM12878 | chr10 | 100,000 | 361,011 | 91,350 | 84,114 |
| Lieberman.GM12878 | chr10 | 250,000 | 129,813 | 96,705 | 83,458 |
| Lieberman.GM12878 | chr10 | 500,000 | 35,256 | 34,572 | 33,536 |
| Lieberman.GM12878 | chr10 | 1,000,000 | 9,001 | 8,853 | 8,647 |
| Lieberman.GM12878 | chr10 | 2,500,000 | 1,540 | 1,488 | 1,439 |
| Lieberman.GM12878 | chr11 | 5,000 | 484,598 | 118,233 | 109,147 |
| Lieberman.GM12878 | chr11 | 10,000 | 469,228 | 131,318 | 124,348 |
| Lieberman.GM12878 | chr11 | 25,000 | 328,871 | 101,285 | 98,269 |
| Lieberman.GM12878 | chr11 | 50,000 | 335,391 | 72,504 | 70,876 |
| Lieberman.GM12878 | chr11 | 100,000 | 372,469 | 100,819 | 92,610 |
| Lieberman.GM12878 | chr11 | 250,000 | 131,103 | 95,125 | 81,196 |
| Lieberman.GM12878 | chr11 | 500,000 | 35,087 | 34,344 | 33,245 |
| Lieberman.GM12878 | chr11 | 1,000,000 | 8,909 | 8,890 | 8,860 |
| Lieberman.GM12878 | chr11 | 2,500,000 | 1,485 | 1,466 | 1,451 |
| Lieberman.GM12878 | chr12 | 5,000 | 452,496 | 119,260 | 34,355 |
| Lieberman.GM12878 | chr12 | 10,000 | 459,416 | 126,364 | 119,449 |
| Lieberman.GM12878 | chr12 | 25,000 | 326,782 | 162,196 | 109,901 |
| Lieberman.GM12878 | chr12 | 50,000 | 275,015 | 116,548 | 75,840 |
| Lieberman.GM12878 | chr12 | 100,000 | 330,069 | 91,506 | 28,598 |
| Lieberman.GM12878 | chr12 | 250,000 | 129,056 | 91,009 | 84,005 |
| Lieberman.GM12878 | chr12 | 500,000 | 34,631 | 33,567 | 32,198 |
| Lieberman.GM12878 | chr12 | 1,000,000 | 8,771 | 8,723 | 8,680 |
| Lieberman.GM12878 | chr12 | 2,500,000 | 1,431 | 1,377 | 1,331 |
| Lieberman.GM12878 | chr13 | 5,000 | 235,766 | 66,668 | 18,877 |
| Lieberman.GM12878 | chr13 | 10,000 | 275,004 | 67,092 | 63,520 |
| Lieberman.GM12878 | chr13 | 25,000 | 210,669 | 65,111 | 63,015 |
| Lieberman.GM12878 | chr13 | 50,000 | 171,677 | 76,351 | 48,043 |
| Lieberman.GM12878 | chr13 | 100,000 | 230,125 | 40,077 | 38,787 |
| Lieberman.GM12878 | chr13 | 250,000 | 73,548 | 67,556 | 55,952 |
| Lieberman.GM12878 | chr13 | 500,000 | 18,714 | 18,630 | 18,494 |
| Lieberman.GM12878 | chr13 | 1,000,000 | 4,753 | 4,710 | 4,652 |
| Lieberman.GM12878 | chr13 | 2,500,000 | 820 | 806 | 794 |
| Lieberman.GM12878 | chr14 | 5,000 | 302,032 | 73,203 | 67,946 |
| Lieberman.GM12878 | chr14 | 10,000 | 301,955 | 82,655 | 78,171 |
| Lieberman.GM12878 | chr14 | 25,000 | 212,484 | 104,436 | 71,493 |
| Lieberman.GM12878 | chr14 | 50,000 | 192,226 | 45,670 | 44,692 |
| Lieberman.GM12878 | chr14 | 100,000 | 225,171 | 48,219 | 45,773 |
| Lieberman.GM12878 | chr14 | 250,000 | 59,555 | 55,615 | 50,563 |
| Lieberman.GM12878 | chr14 | 500,000 | 15,534 | 15,456 | 15,332 |
| Lieberman.GM12878 | chr14 | 1,000,000 | 4,000 | 3,976 | 3,951 |
| Lieberman.GM12878 | chr14 | 2,500,000 | 666 | 666 | 666 |
| Lieberman.GM12878 | chr15 | 5,000 | 307,861 | 79,133 | 22,267 |

*Continued on next page*

Table S7 – *Continued from previous page*

| Dataset identifier | Chr. | Resolution | Non-empty blocks | IDR < 0.05 | IDR < 0.01 |
|---|---|---|---|---|---|
| Lieberman.GM12878 | chr15 | 10,000 | 298,024 | 84,856 | 79,979 |
| Lieberman.GM12878 | chr15 | 25,000 | 200,565 | 100,023 | 68,502 |
| Lieberman.GM12878 | chr15 | 50,000 | 182,864 | 44,409 | 43,482 |
| Lieberman.GM12878 | chr15 | 100,000 | 191,179 | 47,234 | 44,480 |
| Lieberman.GM12878 | chr15 | 250,000 | 49,764 | 45,341 | 41,843 |
| Lieberman.GM12878 | chr15 | 500,000 | 13,501 | 13,381 | 13,198 |
| Lieberman.GM12878 | chr15 | 1,000,000 | 3,486 | 3,474 | 3,460 |
| Lieberman.GM12878 | chr15 | 2,500,000 | 562 | 562 | 562 |
| Lieberman.GM12878 | chr16 | 5,000 | 294,568 | 73,909 | 66,980 |
| Lieberman.GM12878 | chr16 | 10,000 | 284,985 | 77,377 | 72,133 |
| Lieberman.GM12878 | chr16 | 25,000 | 191,094 | 64,648 | 61,992 |
| Lieberman.GM12878 | chr16 | 50,000 | 194,390 | 41,382 | 40,415 |
| Lieberman.GM12878 | chr16 | 100,000 | 175,027 | 55,181 | 50,542 |
| Lieberman.GM12878 | chr16 | 250,000 | 46,386 | 41,366 | 38,288 |
| Lieberman.GM12878 | chr16 | 500,000 | 12,858 | 12,737 | 12,603 |
| Lieberman.GM12878 | chr16 | 1,000,000 | 3,320 | 3,320 | 3,320 |
| Lieberman.GM12878 | chr16 | 2,500,000 | 594 | 589 | 584 |
| Lieberman.GM12878 | chr17 | 5,000 | 361,865 | 92,340 | 83,640 |
| Lieberman.GM12878 | chr17 | 10,000 | 313,381 | 93,428 | 88,389 |
| Lieberman.GM12878 | chr17 | 25,000 | 188,238 | 98,697 | 77,844 |
| Lieberman.GM12878 | chr17 | 50,000 | 170,105 | 39,836 | 38,882 |
| Lieberman.GM12878 | chr17 | 100,000 | 180,407 | 58,239 | 18,133 |
| Lieberman.GM12878 | chr17 | 250,000 | 46,859 | 43,044 | 39,512 |
| Lieberman.GM12878 | chr17 | 500,000 | 12,535 | 12,468 | 12,363 |
| Lieberman.GM12878 | chr17 | 1,000,000 | 3,239 | 3,226 | 3,212 |
| Lieberman.GM12878 | chr17 | 2,500,000 | 528 | 522 | 514 |
| Lieberman.GM12878 | chr18 | 5,000 | 211,392 | 47,932 | 45,556 |
| Lieberman.GM12878 | chr18 | 10,000 | 232,114 | 82,413 | 27,937 |
| Lieberman.GM12878 | chr18 | 25,000 | 173,351 | 90,334 | 63,470 |
| Lieberman.GM12878 | chr18 | 50,000 | 162,375 | 37,368 | 36,715 |
| Lieberman.GM12878 | chr18 | 100,000 | 203,727 | 49,341 | 45,012 |
| Lieberman.GM12878 | chr18 | 250,000 | 44,688 | 43,959 | 42,626 |
| Lieberman.GM12878 | chr18 | 500,000 | 11,435 | 11,350 | 11,208 |
| Lieberman.GM12878 | chr18 | 1,000,000 | 3,001 | 2,981 | 2,960 |
| Lieberman.GM12878 | chr18 | 2,500,000 | 528 | 528 | 528 |
| Lieberman.GM12878 | chr19 | 5,000 | 262,686 | 68,597 | 61,337 |
| Lieberman.GM12878 | chr19 | 10,000 | 235,823 | 69,148 | 65,221 |
| Lieberman.GM12878 | chr19 | 25,000 | 151,773 | 48,195 | 46,620 |
| Lieberman.GM12878 | chr19 | 50,000 | 165,471 | 32,348 | 31,441 |
| Lieberman.GM12878 | chr19 | 100,000 | 114,025 | 68,806 | 38,232 |
| Lieberman.GM12878 | chr19 | 250,000 | 24,810 | 23,787 | 22,410 |
| Lieberman.GM12878 | chr19 | 500,000 | 6,508 | 6,508 | 6,508 |

*Continued on next page*

Table S7 – *Continued from previous page*

| Dataset identifier | Chr. | Resolution | Non-empty blocks | IDR < 0.05 | IDR < 0.01 |
|---|---|---|---|---|---|
| Lieberman.GM12878 | chr19 | 1,000,000 | 1,710 | 1,710 | 1,710 |
| Lieberman.GM12878 | chr19 | 2,500,000 | 276 | 276 | 276 |
| Lieberman.GM12878 | chr20 | 5,000 | 248,229 | 61,439 | 55,749 |
| Lieberman.GM12878 | chr20 | 10,000 | 250,620 | 69,149 | 64,949 |
| Lieberman.GM12878 | chr20 | 25,000 | 168,384 | 89,269 | 68,188 |
| Lieberman.GM12878 | chr20 | 50,000 | 173,955 | 34,616 | 33,939 |
| Lieberman.GM12878 | chr20 | 100,000 | 137,648 | 74,458 | 36,295 |
| Lieberman.GM12878 | chr20 | 250,000 | 28,789 | 28,035 | 26,798 |
| Lieberman.GM12878 | chr20 | 500,000 | 7,361 | 7,353 | 7,342 |
| Lieberman.GM12878 | chr20 | 1,000,000 | 1,891 | 1,891 | 1,891 |
| Lieberman.GM12878 | chr20 | 2,500,000 | 351 | 351 | 351 |
| Lieberman.GM12878 | chr21 | 5,000 | 103,984 | 26,238 | 6,731 |
| Lieberman.GM12878 | chr21 | 10,000 | 106,562 | 29,231 | 27,199 |
| Lieberman.GM12878 | chr21 | 25,000 | 82,123 | 24,613 | 23,732 |
| Lieberman.GM12878 | chr21 | 50,000 | 78,571 | 18,319 | 17,822 |
| Lieberman.GM12878 | chr21 | 100,000 | 44,468 | 23,378 | 21,431 |
| Lieberman.GM12878 | chr21 | 250,000 | 9,486 | 8,682 | 8,263 |
| Lieberman.GM12878 | chr21 | 500,000 | 2,616 | 2,523 | 2,466 |
| Lieberman.GM12878 | chr21 | 1,000,000 | 736 | 728 | 720 |
| Lieberman.GM12878 | chr21 | 2,500,000 | 153 | 152 | 149 |
| Lieberman.GM12878 | chr22 | 5,000 | 172,383 | 45,074 | 39,721 |
| Lieberman.GM12878 | chr22 | 10,000 | 158,446 | 45,952 | 43,071 |
| Lieberman.GM12878 | chr22 | 25,000 | 89,777 | 50,520 | 39,775 |
| Lieberman.GM12878 | chr22 | 50,000 | 100,092 | 18,609 | 18,340 |
| Lieberman.GM12878 | chr22 | 100,000 | 49,585 | 37,930 | 31,245 |
| Lieberman.GM12878 | chr22 | 250,000 | 9,333 | 9,219 | 9,056 |
| Lieberman.GM12878 | chr22 | 500,000 | 2,472 | 2,460 | 2,440 |
| Lieberman.GM12878 | chr22 | 1,000,000 | 661 | 661 | 661 |
| Lieberman.GM12878 | chr22 | 2,500,000 | 120 | 120 | 120 |
| Lieberman.GM12878 | chrX | 5,000 | 269,579 | 65,015 | 63,265 |
| Lieberman.GM12878 | chrX | 10,000 | 329,318 | 73,531 | 70,353 |
| Lieberman.GM12878 | chrX | 25,000 | 314,993 | 86,527 | 83,587 |
| Lieberman.GM12878 | chrX | 50,000 | 333,079 | 110,587 | 44,141 |
| Lieberman.GM12878 | chrX | 100,000 | 440,210 | 132,882 | 39,885 |
| Lieberman.GM12878 | chrX | 250,000 | 171,734 | 103,803 | 98,227 |
| Lieberman.GM12878 | chrX | 500,000 | 46,687 | 40,843 | 36,961 |
| Lieberman.GM12878 | chrX | 1,000,000 | 11,935 | 11,543 | 11,174 |
| Lieberman.GM12878 | chrX | 2,500,000 | 2,016 | 1,954 | 1,909 |
| Lieberman.Patski | chr1 | 5,000 | 32,238 | 128 | 3 |
| Lieberman.Patski | chr1 | 10,000 | 49,770 | 3,739 | 346 |
| Lieberman.Patski | chr1 | 25,000 | 57,694 | 14,445 | 2,139 |
| Lieberman.Patski | chr1 | 50,000 | 46,985 | 16,202 | 13,739 |

*Continued on next page*

Table S7 – *Continued from previous page*

| Dataset identifier | Chr. | Resolution | Non-empty blocks | IDR < 0.05 | IDR < 0.01 |
|---|---|---|---|---|---|
| Lieberman.Patski | chr1 | 100,000 | 31,015 | 11,058 | 10,193 |
| Lieberman.Patski | chr1 | 250,000 | 37,635 | 8,026 | 7,164 |
| Lieberman.Patski | chr1 | 500,000 | 45,687 | 13,811 | 9,553 |
| Lieberman.Patski | chr1 | 1,000,000 | 17,880 | 15,392 | 10,708 |
| Lieberman.Patski | chr1 | 2,500,000 | 3,081 | 3,064 | 3,025 |
| Lieberman.Patski | chr2 | 5,000 | 32,428 | 133 | 3 |
| Lieberman.Patski | chr2 | 10,000 | 49,666 | 4,990 | 495 |
| Lieberman.Patski | chr2 | 25,000 | 55,247 | 19,390 | 15,514 |
| Lieberman.Patski | chr2 | 50,000 | 44,683 | 15,333 | 13,364 |
| Lieberman.Patski | chr2 | 100,000 | 28,313 | 13,710 | 5,902 |
| Lieberman.Patski | chr2 | 250,000 | 38,246 | 7,093 | 6,404 |
| Lieberman.Patski | chr2 | 500,000 | 39,223 | 14,110 | 10,364 |
| Lieberman.Patski | chr2 | 1,000,000 | 15,077 | 12,498 | 9,471 |
| Lieberman.Patski | chr2 | 2,500,000 | 2,597 | 2,594 | 2,590 |
| Lieberman.Patski | chr3 | 5,000 | 52,319 | 4,386 | 935 |
| Lieberman.Patski | chr3 | 10,000 | 67,860 | 18,686 | 11,654 |
| Lieberman.Patski | chr3 | 25,000 | 73,569 | 22,960 | 17,814 |
| Lieberman.Patski | chr3 | 50,000 | 57,542 | 19,022 | 16,067 |
| Lieberman.Patski | chr3 | 100,000 | 39,804 | 15,533 | 4,614 |
| Lieberman.Patski | chr3 | 250,000 | 72,703 | 12,967 | 10,383 |
| Lieberman.Patski | chr3 | 500,000 | 43,092 | 24,836 | 9,690 |
| Lieberman.Patski | chr3 | 1,000,000 | 12,402 | 12,230 | 11,715 |
| Lieberman.Patski | chr3 | 2,500,000 | 2,016 | 2,016 | 2,016 |
| Lieberman.Patski | chr4 | 5,000 | 14,719 | 42 | 0 |
| Lieberman.Patski | chr4 | 10,000 | 27,131 | 0 | 0 |
| Lieberman.Patski | chr4 | 25,000 | 31,165 | 3,260 | 194 |
| Lieberman.Patski | chr4 | 50,000 | 26,064 | 7,200 | 921 |
| Lieberman.Patski | chr4 | 100,000 | 17,867 | 6,860 | 5,696 |
| Lieberman.Patski | chr4 | 250,000 | 16,473 | 4,035 | 3,619 |
| Lieberman.Patski | chr4 | 500,000 | 24,853 | 4,569 | 3,303 |
| Lieberman.Patski | chr4 | 1,000,000 | 10,507 | 8,322 | 5,023 |
| Lieberman.Patski | chr4 | 2,500,000 | 1,953 | 1,941 | 1,900 |
| Lieberman.Patski | chr5 | 5,000 | 40,329 | 3,891 | 1,256 |
| Lieberman.Patski | chr5 | 10,000 | 52,515 | 17,060 | 11,445 |
| Lieberman.Patski | chr5 | 25,000 | 54,896 | 18,485 | 15,488 |
| Lieberman.Patski | chr5 | 50,000 | 42,097 | 16,245 | 4,385 |
| Lieberman.Patski | chr5 | 100,000 | 29,876 | 9,495 | 8,930 |
| Lieberman.Patski | chr5 | 250,000 | 51,853 | 9,891 | 8,256 |
| Lieberman.Patski | chr5 | 500,000 | 35,429 | 18,231 | 6,343 |
| Lieberman.Patski | chr5 | 1,000,000 | 10,849 | 10,136 | 7,696 |
| Lieberman.Patski | chr5 | 2,500,000 | 1,830 | 1,774 | 1,638 |
| Lieberman.Patski | chr6 | 5,000 | 25,092 | 120 | 1 |

*Continued on next page*

Table S7 – *Continued from previous page*

| Dataset identifier | Chr. | Resolution | Non-empty blocks | IDR < 0.05 | IDR < 0.01 |
|---|---|---|---|---|---|
| Lieberman.Patski | chr6 | 10,000 | 38,470 | 2,921 | 300 |
| Lieberman.Patski | chr6 | 25,000 | 42,422 | 10,235 | 1,500 |
| Lieberman.Patski | chr6 | 50,000 | 34,661 | 11,663 | 9,718 |
| Lieberman.Patski | chr6 | 100,000 | 24,058 | 8,348 | 7,279 |
| Lieberman.Patski | chr6 | 250,000 | 34,218 | 6,549 | 5,558 |
| Lieberman.Patski | chr6 | 500,000 | 31,216 | 8,700 | 2,361 |
| Lieberman.Patski | chr6 | 1,000,000 | 10,738 | 9,562 | 5,962 |
| Lieberman.Patski | chr6 | 2,500,000 | 1,770 | 1,753 | 1,698 |
| Lieberman.Patski | chr7 | 5,000 | 24,110 | 72 | 0 |
| Lieberman.Patski | chr7 | 10,000 | 35,316 | 9,224 | 4,988 |
| Lieberman.Patski | chr7 | 25,000 | 38,416 | 9,125 | 1,184 |
| Lieberman.Patski | chr7 | 50,000 | 29,309 | 10,484 | 8,740 |
| Lieberman.Patski | chr7 | 100,000 | 19,494 | 7,159 | 6,456 |
| Lieberman.Patski | chr7 | 250,000 | 26,871 | 5,339 | 4,483 |
| Lieberman.Patski | chr7 | 500,000 | 27,236 | 7,940 | 5,585 |
| Lieberman.Patski | chr7 | 1,000,000 | 9,250 | 8,255 | 4,974 |
| Lieberman.Patski | chr7 | 2,500,000 | 1,658 | 1,641 | 1,607 |
| Lieberman.Patski | chr8 | 5,000 | 24,470 | 148 | 8 |
| Lieberman.Patski | chr8 | 10,000 | 35,669 | 9,099 | 4,823 |
| Lieberman.Patski | chr8 | 25,000 | 39,989 | 9,814 | 1,466 |
| Lieberman.Patski | chr8 | 50,000 | 32,193 | 11,415 | 9,705 |
| Lieberman.Patski | chr8 | 100,000 | 21,247 | 7,339 | 6,681 |
| Lieberman.Patski | chr8 | 250,000 | 28,262 | 5,479 | 4,885 |
| Lieberman.Patski | chr8 | 500,000 | 25,255 | 8,088 | 2,332 |
| Lieberman.Patski | chr8 | 1,000,000 | 7,896 | 7,788 | 7,440 |
| Lieberman.Patski | chr8 | 2,500,000 | 1,326 | 1,326 | 1,326 |
| Lieberman.Patski | chr9 | 5,000 | 25,348 | 96 | 4 |
| Lieberman.Patski | chr9 | 10,000 | 37,108 | 11,317 | 6,311 |
| Lieberman.Patski | chr9 | 25,000 | 42,443 | 11,559 | 1,691 |
| Lieberman.Patski | chr9 | 50,000 | 32,181 | 11,760 | 10,179 |
| Lieberman.Patski | chr9 | 100,000 | 20,535 | 9,590 | 4,223 |
| Lieberman.Patski | chr9 | 250,000 | 25,596 | 4,655 | 4,359 |
| Lieberman.Patski | chr9 | 500,000 | 23,946 | 9,745 | 2,988 |
| Lieberman.Patski | chr9 | 1,000,000 | 7,458 | 7,202 | 6,490 |
| Lieberman.Patski | chr9 | 2,500,000 | 1,225 | 1,193 | 1,116 |
| Lieberman.Patski | chr10 | 5,000 | 22,338 | 104 | 2 |
| Lieberman.Patski | chr10 | 10,000 | 34,309 | 4,013 | 409 |
| Lieberman.Patski | chr10 | 25,000 | 40,844 | 11,524 | 1,861 |
| Lieberman.Patski | chr10 | 50,000 | 33,877 | 11,382 | 10,206 |
| Lieberman.Patski | chr10 | 100,000 | 21,891 | 10,667 | 4,784 |
| Lieberman.Patski | chr10 | 250,000 | 25,870 | 4,527 | 4,347 |
| Lieberman.Patski | chr10 | 500,000 | 25,701 | 8,273 | 2,359 |

*Continued on next page*

Table S7 – *Continued from previous page*

| Dataset identifier | Chr. | Resolution | Non-empty blocks | IDR < 0.05 | IDR < 0.01 |
|---|---|---|---|---|---|
| Lieberman.Patski | chr10 | 1,000,000 | 8,224 | 8,016 | 7,386 |
| Lieberman.Patski | chr10 | 2,500,000 | 1,378 | 1,358 | 1,311 |
| Lieberman.Patski | chr11 | 5,000 | 27,350 | 66 | 0 |
| Lieberman.Patski | chr11 | 10,000 | 39,517 | 3,920 | 312 |
| Lieberman.Patski | chr11 | 25,000 | 40,085 | 14,247 | 11,471 |
| Lieberman.Patski | chr11 | 50,000 | 29,454 | 11,250 | 9,906 |
| Lieberman.Patski | chr11 | 100,000 | 19,059 | 9,181 | 3,706 |
| Lieberman.Patski | chr11 | 250,000 | 29,183 | 5,122 | 4,549 |
| Lieberman.Patski | chr11 | 500,000 | 20,714 | 10,762 | 3,743 |
| Lieberman.Patski | chr11 | 1,000,000 | 6,836 | 6,388 | 5,311 |
| Lieberman.Patski | chr11 | 2,500,000 | 1,176 | 1,176 | 1,176 |
| Lieberman.Patski | chr12 | 5,000 | 13,092 | 0 | 0 |
| Lieberman.Patski | chr12 | 10,000 | 24,422 | 423 | 16 |
| Lieberman.Patski | chr12 | 25,000 | 28,186 | 8,915 | 6,257 |
| Lieberman.Patski | chr12 | 50,000 | 25,070 | 6,416 | 959 |
| Lieberman.Patski | chr12 | 100,000 | 16,952 | 6,357 | 1,460 |
| Lieberman.Patski | chr12 | 250,000 | 17,503 | 2,509 | 494 |
| Lieberman.Patski | chr12 | 500,000 | 19,610 | 2,863 | 708 |
| Lieberman.Patski | chr12 | 1,000,000 | 6,510 | 4,679 | 1,424 |
| Lieberman.Patski | chr12 | 2,500,000 | 1,136 | 1,136 | 1,136 |
| Lieberman.Patski | chr13 | 5,000 | 20,026 | 191 | 0 |
| Lieberman.Patski | chr13 | 10,000 | 30,135 | 8,089 | 3,609 |
| Lieberman.Patski | chr13 | 25,000 | 37,466 | 11,973 | 9,511 |
| Lieberman.Patski | chr13 | 50,000 | 30,298 | 10,864 | 9,499 |
| Lieberman.Patski | chr13 | 100,000 | 19,848 | 8,989 | 3,354 |
| Lieberman.Patski | chr13 | 250,000 | 20,516 | 4,323 | 4,065 |
| Lieberman.Patski | chr13 | 500,000 | 20,506 | 6,555 | 1,835 |
| Lieberman.Patski | chr13 | 1,000,000 | 6,899 | 5,988 | 4,304 |
| Lieberman.Patski | chr13 | 2,500,000 | 1,169 | 1,155 | 1,123 |
| Lieberman.Patski | chr14 | 5,000 | 16,664 | 0 | 0 |
| Lieberman.Patski | chr14 | 10,000 | 26,991 | 1,275 | 91 |
| Lieberman.Patski | chr14 | 25,000 | 31,693 | 9,960 | 7,529 |
| Lieberman.Patski | chr14 | 50,000 | 25,932 | 7,776 | 1,324 |
| Lieberman.Patski | chr14 | 100,000 | 17,567 | 5,892 | 5,198 |
| Lieberman.Patski | chr14 | 250,000 | 21,956 | 4,491 | 3,934 |
| Lieberman.Patski | chr14 | 500,000 | 21,556 | 5,393 | 1,515 |
| Lieberman.Patski | chr14 | 1,000,000 | 6,908 | 6,571 | 5,166 |
| Lieberman.Patski | chr14 | 2,500,000 | 1,197 | 1,178 | 1,153 |
| Lieberman.Patski | chr15 | 5,000 | 20,456 | 95 | 1 |
| Lieberman.Patski | chr15 | 10,000 | 30,156 | 7,855 | 4,317 |
| Lieberman.Patski | chr15 | 25,000 | 33,659 | 11,134 | 8,925 |
| Lieberman.Patski | chr15 | 50,000 | 26,573 | 9,281 | 7,963 |

*Continued on next page*

Table S7 – *Continued from previous page*

| Dataset identifier | Chr. | Resolution | Non-empty blocks | IDR < 0.05 | IDR < 0.01 |
|---|---|---|---|---|---|
| Lieberman.Patski | chr15 | 100,000 | 17,898 | 6,117 | 5,752 |
| Lieberman.Patski | chr15 | 250,000 | 24,739 | 4,884 | 4,334 |
| Lieberman.Patski | chr15 | 500,000 | 17,505 | 6,958 | 2,231 |
| Lieberman.Patski | chr15 | 1,000,000 | 5,090 | 4,765 | 3,294 |
| Lieberman.Patski | chr15 | 2,500,000 | 861 | 859 | 852 |
| Lieberman.Patski | chr16 | 5,000 | 15,798 | 55 | 0 |
| Lieberman.Patski | chr16 | 10,000 | 24,796 | 6,322 | 3,392 |
| Lieberman.Patski | chr16 | 25,000 | 28,585 | 9,955 | 7,757 |
| Lieberman.Patski | chr16 | 50,000 | 23,335 | 8,218 | 7,321 |
| Lieberman.Patski | chr16 | 100,000 | 16,185 | 5,383 | 5,074 |
| Lieberman.Patski | chr16 | 250,000 | 22,867 | 4,020 | 3,731 |
| Lieberman.Patski | chr16 | 500,000 | 15,875 | 9,069 | 3,542 |
| Lieberman.Patski | chr16 | 1,000,000 | 4,575 | 4,336 | 3,504 |
| Lieberman.Patski | chr16 | 2,500,000 | 780 | 772 | 757 |
| Lieberman.Patski | chr17 | 5,000 | 17,464 | 29 | 0 |
| Lieberman.Patski | chr17 | 10,000 | 25,690 | 2,435 | 154 |
| Lieberman.Patski | chr17 | 25,000 | 27,525 | 9,512 | 7,301 |
| Lieberman.Patski | chr17 | 50,000 | 22,758 | 7,580 | 1,501 |
| Lieberman.Patski | chr17 | 100,000 | 14,732 | 5,309 | 4,719 |
| Lieberman.Patski | chr17 | 250,000 | 20,904 | 3,723 | 3,067 |
| Lieberman.Patski | chr17 | 500,000 | 14,104 | 4,926 | 1,374 |
| Lieberman.Patski | chr17 | 1,000,000 | 4,247 | 3,661 | 965 |
| Lieberman.Patski | chr17 | 2,500,000 | 703 | 638 | 429 |
| Lieberman.Patski | chr18 | 5,000 | 15,706 | 77 | 6 |
| Lieberman.Patski | chr18 | 10,000 | 24,024 | 6,515 | 3,560 |
| Lieberman.Patski | chr18 | 25,000 | 28,112 | 9,656 | 7,793 |
| Lieberman.Patski | chr18 | 50,000 | 23,350 | 8,009 | 6,986 |
| Lieberman.Patski | chr18 | 100,000 | 14,953 | 7,321 | 3,227 |
| Lieberman.Patski | chr18 | 250,000 | 22,521 | 4,087 | 3,649 |
| Lieberman.Patski | chr18 | 500,000 | 14,287 | 8,634 | 3,658 |
| Lieberman.Patski | chr18 | 1,000,000 | 3,915 | 3,906 | 3,886 |
| Lieberman.Patski | chr18 | 2,500,000 | 666 | 656 | 631 |
| Lieberman.Patski | chr19 | 5,000 | 14,235 | 91 | 4 |
| Lieberman.Patski | chr19 | 10,000 | 19,920 | 6,578 | 3,939 |
| Lieberman.Patski | chr19 | 25,000 | 21,167 | 7,232 | 6,023 |
| Lieberman.Patski | chr19 | 50,000 | 15,252 | 5,691 | 5,115 |
| Lieberman.Patski | chr19 | 100,000 | 9,893 | 4,744 | 2,478 |
| Lieberman.Patski | chr19 | 250,000 | 15,083 | 2,778 | 2,348 |
| Lieberman.Patski | chr19 | 500,000 | 6,585 | 3,586 | 1,417 |
| Lieberman.Patski | chr19 | 1,000,000 | 1,768 | 1,368 | 407 |
| Lieberman.Patski | chr19 | 2,500,000 | 300 | 286 | 244 |
| Lieberman.Patski | chrX | 5,000 | 9,548 | 0 | 0 |

*Continued on next page*

Table S7 – *Continued from previous page*

| Dataset identifier | Chr. | Resolution | Non-empty blocks | IDR < 0.05 | IDR < 0.01 |
|---|---|---|---|---|---|
| Lieberman.Patski | chrX | 10,000 | 20,875 | 506 | 47 |
| Lieberman.Patski | chrX | 25,000 | 21,528 | 2,182 | 278 |
| Lieberman.Patski | chrX | 50,000 | 21,134 | 6,041 | 4,532 |
| Lieberman.Patski | chrX | 100,000 | 18,567 | 6,090 | 4,840 |
| Lieberman.Patski | chrX | 250,000 | 26,115 | 5,461 | 4,533 |
| Lieberman.Patski | chrX | 500,000 | 37,066 | 4,580 | 3,305 |
| Lieberman.Patski | chrX | 1,000,000 | 11,361 | 2,002 | 1,216 |
| Lieberman.Patski | chrX | 2,500,000 | 2,147 | 1,889 | 498 |
| Skok.NSD2 | chr1 | 5,000 | 7,183 | 324 | 146 |
| Skok.NSD2 | chr1 | 10,000 | 59,428 | 2,586 | 422 |
| Skok.NSD2 | chr1 | 25,000 | 104,544 | 34,476 | 28,899 |
| Skok.NSD2 | chr1 | 50,000 | 85,480 | 27,070 | 25,441 |
| Skok.NSD2 | chr1 | 100,000 | 57,925 | 26,805 | 15,134 |
| Skok.NSD2 | chr1 | 250,000 | 56,502 | 10,362 | 2,946 |
| Skok.NSD2 | chr1 | 500,000 | 35,766 | 14,086 | 11,650 |
| Skok.NSD2 | chr1 | 1,000,000 | 16,333 | 9,140 | 8,741 |
| Skok.NSD2 | chr1 | 2,500,000 | 4,382 | 3,290 | 2,540 |
| Skok.NSD2 | chr2 | 5,000 | 369 | 10 | 0 |
| Skok.NSD2 | chr2 | 10,000 | 36,839 | 1,478 | 659 |
| Skok.NSD2 | chr2 | 25,000 | 98,219 | 33,740 | 28,302 |
| Skok.NSD2 | chr2 | 50,000 | 91,055 | 27,868 | 25,774 |
| Skok.NSD2 | chr2 | 100,000 | 57,322 | 30,194 | 21,454 |
| Skok.NSD2 | chr2 | 250,000 | 49,081 | 9,775 | 9,458 |
| Skok.NSD2 | chr2 | 500,000 | 47,012 | 14,497 | 11,524 |
| Skok.NSD2 | chr2 | 1,000,000 | 24,521 | 15,432 | 11,299 |
| Skok.NSD2 | chr2 | 2,500,000 | 4,753 | 4,446 | 3,917 |
| Skok.NSD2 | chr3 | 5,000 | 189 | 109 | 0 |
| Skok.NSD2 | chr3 | 10,000 | 24,488 | 681 | 284 |
| Skok.NSD2 | chr3 | 25,000 | 73,880 | 19,835 | 3,508 |
| Skok.NSD2 | chr3 | 50,000 | 70,204 | 22,088 | 20,317 |
| Skok.NSD2 | chr3 | 100,000 | 45,350 | 22,624 | 13,028 |
| Skok.NSD2 | chr3 | 250,000 | 38,716 | 7,441 | 7,252 |
| Skok.NSD2 | chr3 | 500,000 | 33,083 | 10,380 | 8,102 |
| Skok.NSD2 | chr3 | 1,000,000 | 16,338 | 8,794 | 7,981 |
| Skok.NSD2 | chr3 | 2,500,000 | 3,231 | 2,387 | 2,142 |
| Skok.NSD2 | chr4 | 5,000 | 147 | 124 | 19 |
| Skok.NSD2 | chr4 | 10,000 | 27,788 | 832 | 378 |
| Skok.NSD2 | chr4 | 25,000 | 78,558 | 26,161 | 21,253 |
| Skok.NSD2 | chr4 | 50,000 | 77,172 | 22,768 | 20,844 |
| Skok.NSD2 | chr4 | 100,000 | 49,559 | 17,749 | 16,812 |
| Skok.NSD2 | chr4 | 250,000 | 44,461 | 8,918 | 8,594 |
| Skok.NSD2 | chr4 | 500,000 | 36,151 | 11,155 | 3,227 |

*Continued on next page*

Table S7 – *Continued from previous page*

| Dataset identifier | Chr. | Resolution | Non-empty blocks | IDR < 0.05 | IDR < 0.01 |
|---|---|---|---|---|---|
| Skok.NSD2 | chr4 | 1,000,000 | 14,713 | 10,762 | 8,991 |
| Skok.NSD2 | chr4 | 2,500,000 | 2,946 | 2,566 | 2,222 |
| Skok.NSD2 | chr5 | 5,000 | 185 | 0 | 0 |
| Skok.NSD2 | chr5 | 10,000 | 25,695 | 684 | 307 |
| Skok.NSD2 | chr5 | 25,000 | 74,824 | 25,411 | 20,550 |
| Skok.NSD2 | chr5 | 50,000 | 71,639 | 23,117 | 20,737 |
| Skok.NSD2 | chr5 | 100,000 | 45,220 | 24,230 | 17,046 |
| Skok.NSD2 | chr5 | 250,000 | 35,620 | 7,628 | 7,413 |
| Skok.NSD2 | chr5 | 500,000 | 32,006 | 9,835 | 7,855 |
| Skok.NSD2 | chr5 | 1,000,000 | 14,541 | 9,831 | 7,296 |
| Skok.NSD2 | chr5 | 2,500,000 | 2,640 | 2,529 | 2,322 |
| Skok.NSD2 | chr6 | 5,000 | 57 | 52 | 47 |
| Skok.NSD2 | chr6 | 10,000 | 13,274 | 602 | 380 |
| Skok.NSD2 | chr6 | 25,000 | 53,999 | 19,303 | 14,980 |
| Skok.NSD2 | chr6 | 50,000 | 55,872 | 16,844 | 15,421 |
| Skok.NSD2 | chr6 | 100,000 | 35,996 | 12,697 | 12,029 |
| Skok.NSD2 | chr6 | 250,000 | 26,805 | 6,028 | 5,886 |
| Skok.NSD2 | chr6 | 500,000 | 23,632 | 6,469 | 5,457 |
| Skok.NSD2 | chr6 | 1,000,000 | 11,825 | 6,490 | 6,062 |
| Skok.NSD2 | chr6 | 2,500,000 | 2,414 | 2,090 | 1,596 |
| Skok.NSD2 | chr7 | 5,000 | 511 | 21 | 0 |
| Skok.NSD2 | chr7 | 10,000 | 35,545 | 104 | 0 |
| Skok.NSD2 | chr7 | 25,000 | 78,273 | 25,909 | 22,158 |
| Skok.NSD2 | chr7 | 50,000 | 67,147 | 22,188 | 20,377 |
| Skok.NSD2 | chr7 | 100,000 | 41,681 | 21,105 | 13,848 |
| Skok.NSD2 | chr7 | 250,000 | 40,967 | 7,832 | 7,433 |
| Skok.NSD2 | chr7 | 500,000 | 29,883 | 10,522 | 8,104 |
| Skok.NSD2 | chr7 | 1,000,000 | 12,203 | 7,217 | 5,995 |
| Skok.NSD2 | chr7 | 2,500,000 | 2,079 | 1,958 | 1,643 |
| Skok.NSD2 | chr8 | 5,000 | 418 | 4 | 0 |
| Skok.NSD2 | chr8 | 10,000 | 24,591 | 894 | 407 |
| Skok.NSD2 | chr8 | 25,000 | 63,627 | 21,560 | 17,580 |
| Skok.NSD2 | chr8 | 50,000 | 58,076 | 19,876 | 17,854 |
| Skok.NSD2 | chr8 | 100,000 | 36,960 | 13,713 | 12,964 |
| Skok.NSD2 | chr8 | 250,000 | 32,135 | 6,724 | 6,493 |
| Skok.NSD2 | chr8 | 500,000 | 26,902 | 8,680 | 2,510 |
| Skok.NSD2 | chr8 | 1,000,000 | 10,121 | 6,866 | 5,558 |
| Skok.NSD2 | chr8 | 2,500,000 | 1,721 | 1,635 | 1,454 |
| Skok.NSD2 | chr9 | 5,000 | 95 | 4 | 0 |
| Skok.NSD2 | chr9 | 10,000 | 18,667 | 430 | 192 |
| Skok.NSD2 | chr9 | 25,000 | 47,214 | 17,042 | 13,965 |
| Skok.NSD2 | chr9 | 50,000 | 44,072 | 15,894 | 3,803 |

*Continued on next page*

Table S7 – *Continued from previous page*

| Dataset identifier | Chr. | Resolution | Non-empty blocks | IDR < 0.05 | IDR < 0.01 |
|---|---|---|---|---|---|
| Skok.NSD2 | chr9 | 100,000 | 27,379 | 10,560 | 9,873 |
| Skok.NSD2 | chr9 | 250,000 | 22,980 | 4,569 | 4,403 |
| Skok.NSD2 | chr9 | 500,000 | 13,514 | 6,081 | 2,221 |
| Skok.NSD2 | chr9 | 1,000,000 | 6,356 | 4,035 | 2,703 |
| Skok.NSD2 | chr9 | 2,500,000 | 1,252 | 1,162 | 1,008 |
| Skok.NSD2 | chr10 | 5,000 | 95 | 65 | 0 |
| Skok.NSD2 | chr10 | 10,000 | 19,750 | 642 | 337 |
| Skok.NSD2 | chr10 | 25,000 | 54,909 | 15,644 | 2,380 |
| Skok.NSD2 | chr10 | 50,000 | 49,513 | 21,305 | 7,411 |
| Skok.NSD2 | chr10 | 100,000 | 32,416 | 15,834 | 9,450 |
| Skok.NSD2 | chr10 | 250,000 | 27,151 | 5,668 | 5,464 |
| Skok.NSD2 | chr10 | 500,000 | 21,014 | 6,680 | 5,790 |
| Skok.NSD2 | chr10 | 1,000,000 | 8,651 | 5,976 | 4,531 |
| Skok.NSD2 | chr10 | 2,500,000 | 1,485 | 1,334 | 994 |
| Skok.NSD2 | chr11 | 5,000 | 199 | 38 | 0 |
| Skok.NSD2 | chr11 | 10,000 | 15,021 | 496 | 254 |
| Skok.NSD2 | chr11 | 25,000 | 45,399 | 16,522 | 13,331 |
| Skok.NSD2 | chr11 | 50,000 | 42,376 | 16,993 | 5,256 |
| Skok.NSD2 | chr11 | 100,000 | 28,023 | 13,293 | 6,520 |
| Skok.NSD2 | chr11 | 250,000 | 22,134 | 4,763 | 4,604 |
| Skok.NSD2 | chr11 | 500,000 | 16,807 | 4,774 | 4,107 |
| Skok.NSD2 | chr11 | 1,000,000 | 6,159 | 4,933 | 4,177 |
| Skok.NSD2 | chr11 | 2,500,000 | 1,504 | 1,096 | 1,076 |
| Skok.NSD2 | chr12 | 5,000 | 2,079 | 68 | 29 |
| Skok.NSD2 | chr12 | 10,000 | 24,070 | 3,065 | 1,994 |
| Skok.NSD2 | chr12 | 25,000 | 56,341 | 18,753 | 14,732 |
| Skok.NSD2 | chr12 | 50,000 | 49,026 | 18,592 | 5,094 |
| Skok.NSD2 | chr12 | 100,000 | 33,582 | 14,341 | 6,154 |
| Skok.NSD2 | chr12 | 250,000 | 24,672 | 6,201 | 5,817 |
| Skok.NSD2 | chr12 | 500,000 | 20,059 | 4,916 | 1,333 |
| Skok.NSD2 | chr12 | 1,000,000 | 7,375 | 5,353 | 4,045 |
| Skok.NSD2 | chr12 | 2,500,000 | 1,462 | 1,422 | 1,339 |
| Skok.NSD2 | chr13 | 10,000 | 823 | 6 | 0 |
| Skok.NSD2 | chr13 | 25,000 | 20,749 | 4,916 | 526 |
| Skok.NSD2 | chr13 | 50,000 | 25,725 | 8,562 | 7,490 |
| Skok.NSD2 | chr13 | 100,000 | 18,259 | 7,401 | 6,956 |
| Skok.NSD2 | chr13 | 250,000 | 10,826 | 4,734 | 2,606 |
| Skok.NSD2 | chr13 | 500,000 | 11,534 | 2,407 | 2,151 |
| Skok.NSD2 | chr13 | 1,000,000 | 4,629 | 2,628 | 1,233 |
| Skok.NSD2 | chr13 | 2,500,000 | 782 | 665 | 384 |
| Skok.NSD2 | chr14 | 5,000 | 46 | 46 | 46 |
| Skok.NSD2 | chr14 | 10,000 | 6,832 | 162 | 82 |

*Continued on next page*

Table S7 – *Continued from previous page*

| **Dataset identifier** | **Chr.** | **Resolution** | **Non-empty blocks** | **IDR < 0.05** | **IDR < 0.01** |
|---|---|---:|---:|---:|---:|
| Skok.NSD2 | chr14 | 25,000 | 27,449 | 9,924 | 7,765 |
| Skok.NSD2 | chr14 | 50,000 | 26,681 | 8,389 | 7,788 |
| Skok.NSD2 | chr14 | 100,000 | 18,180 | 8,647 | 4,796 |
| Skok.NSD2 | chr14 | 250,000 | 11,792 | 4,404 | 2,216 |
| Skok.NSD2 | chr14 | 500,000 | 11,121 | 2,526 | 2,273 |
| Skok.NSD2 | chr14 | 1,000,000 | 3,781 | 3,116 | 1,835 |
| Skok.NSD2 | chr14 | 2,500,000 | 665 | 637 | 584 |
| Skok.NSD2 | chr15 | 10,000 | 11,170 | 448 | 256 |
| Skok.NSD2 | chr15 | 25,000 | 30,972 | 11,450 | 9,627 |
| Skok.NSD2 | chr15 | 50,000 | 27,323 | 9,318 | 8,682 |
| Skok.NSD2 | chr15 | 100,000 | 17,367 | 8,771 | 5,895 |
| Skok.NSD2 | chr15 | 250,000 | 12,969 | 2,863 | 2,777 |
| Skok.NSD2 | chr15 | 500,000 | 10,173 | 2,556 | 2,240 |
| Skok.NSD2 | chr15 | 1,000,000 | 3,329 | 2,995 | 2,054 |
| Skok.NSD2 | chr15 | 2,500,000 | 574 | 574 | 574 |
| Skok.NSD2 | chr16 | 5,000 | 402 | 25 | 15 |
| Skok.NSD2 | chr16 | 10,000 | 17,962 | 1,069 | 452 |
| Skok.NSD2 | chr16 | 25,000 | 33,544 | 11,167 | 2,069 |
| Skok.NSD2 | chr16 | 50,000 | 28,227 | 12,669 | 4,876 |
| Skok.NSD2 | chr16 | 100,000 | 18,720 | 8,946 | 4,969 |
| Skok.NSD2 | chr16 | 250,000 | 14,443 | 3,629 | 3,419 |
| Skok.NSD2 | chr16 | 500,000 | 9,171 | 2,667 | 2,428 |
| Skok.NSD2 | chr16 | 1,000,000 | 3,184 | 2,597 | 2,115 |
| Skok.NSD2 | chr16 | 2,500,000 | 582 | 580 | 577 |
| Skok.NSD2 | chr17 | 5,000 | 200 | 25 | 0 |
| Skok.NSD2 | chr17 | 10,000 | 15,381 | 586 | 236 |
| Skok.NSD2 | chr17 | 25,000 | 28,274 | 9,523 | 1,440 |
| Skok.NSD2 | chr17 | 50,000 | 22,159 | 10,859 | 3,957 |
| Skok.NSD2 | chr17 | 100,000 | 13,581 | 7,498 | 5,813 |
| Skok.NSD2 | chr17 | 250,000 | 11,035 | 2,133 | 2,064 |
| Skok.NSD2 | chr17 | 500,000 | 8,360 | 2,684 | 2,100 |
| Skok.NSD2 | chr17 | 1,000,000 | 3,031 | 2,437 | 1,603 |
| Skok.NSD2 | chr17 | 2,500,000 | 557 | 557 | 557 |
| Skok.NSD2 | chr18 | 5,000 | 287 | 2 | 0 |
| Skok.NSD2 | chr18 | 10,000 | 15,712 | 921 | 392 |
| Skok.NSD2 | chr18 | 25,000 | 36,877 | 12,341 | 10,286 |
| Skok.NSD2 | chr18 | 50,000 | 32,173 | 10,657 | 9,738 |
| Skok.NSD2 | chr18 | 100,000 | 20,509 | 9,811 | 5,015 |
| Skok.NSD2 | chr18 | 250,000 | 17,938 | 3,904 | 3,694 |
| Skok.NSD2 | chr18 | 500,000 | 10,243 | 3,408 | 2,947 |
| Skok.NSD2 | chr18 | 1,000,000 | 2,807 | 2,586 | 1,691 |
| Skok.NSD2 | chr18 | 2,500,000 | 507 | 493 | 462 |

*Continued on next page*

Table S7 – *Continued from previous page*

| **Dataset identifier** | **Chr.** | **Resolution** | **Non-empty blocks** | **IDR < 0.05** | **IDR < 0.01** |
|---|---|---|---|---|---|
| Skok.NSD2 | chr19 | 5,000 | 277 | 43 | 0 |
| Skok.NSD2 | chr19 | 10,000 | 13,189 | 161 | 12 |
| Skok.NSD2 | chr19 | 25,000 | 24,387 | 8,309 | 1,586 |
| Skok.NSD2 | chr19 | 50,000 | 20,006 | 10,060 | 6,022 |
| Skok.NSD2 | chr19 | 100,000 | 14,277 | 4,358 | 4,233 |
| Skok.NSD2 | chr19 | 250,000 | 12,180 | 2,532 | 2,399 |
| Skok.NSD2 | chr19 | 500,000 | 5,627 | 3,490 | 1,946 |
| Skok.NSD2 | chr19 | 1,000,000 | 1,650 | 1,592 | 1,471 |
| Skok.NSD2 | chr19 | 2,500,000 | 298 | 298 | 298 |
| Skok.NSD2 | chr20 | 5,000 | 267 | 0 | 0 |
| Skok.NSD2 | chr20 | 10,000 | 12,988 | 143 | 12 |
| Skok.NSD2 | chr20 | 25,000 | 26,001 | 8,874 | 7,561 |
| Skok.NSD2 | chr20 | 50,000 | 21,922 | 9,938 | 4,225 |
| Skok.NSD2 | chr20 | 100,000 | 14,783 | 6,789 | 4,033 |
| Skok.NSD2 | chr20 | 250,000 | 10,476 | 2,396 | 2,242 |
| Skok.NSD2 | chr20 | 500,000 | 6,438 | 2,315 | 2,144 |
| Skok.NSD2 | chr20 | 1,000,000 | 2,000 | 1,673 | 1,164 |
| Skok.NSD2 | chr20 | 2,500,000 | 351 | 326 | 268 |
| Skok.NSD2 | chr21 | 5,000 | 56 | 56 | 56 |
| Skok.NSD2 | chr21 | 10,000 | 7,385 | 192 | 78 |
| Skok.NSD2 | chr21 | 25,000 | 16,158 | 5,641 | 4,748 |
| Skok.NSD2 | chr21 | 50,000 | 14,240 | 4,842 | 4,417 |
| Skok.NSD2 | chr21 | 100,000 | 9,131 | 4,749 | 2,951 |
| Skok.NSD2 | chr21 | 250,000 | 6,552 | 1,402 | 1,362 |
| Skok.NSD2 | chr21 | 500,000 | 2,431 | 1,927 | 1,561 |
| Skok.NSD2 | chr21 | 1,000,000 | 699 | 658 | 612 |
| Skok.NSD2 | chr21 | 2,500,000 | 153 | 129 | 122 |
| Skok.NSD2 | chr22 | 5,000 | 26 | 26 | 26 |
| Skok.NSD2 | chr22 | 10,000 | 6,104 | 109 | 41 |
| Skok.NSD2 | chr22 | 25,000 | 13,006 | 5,129 | 4,450 |
| Skok.NSD2 | chr22 | 50,000 | 10,860 | 5,022 | 2,071 |
| Skok.NSD2 | chr22 | 100,000 | 6,471 | 3,523 | 2,817 |
| Skok.NSD2 | chr22 | 250,000 | 5,325 | 1,302 | 1,152 |
| Skok.NSD2 | chr22 | 500,000 | 2,277 | 1,505 | 930 |
| Skok.NSD2 | chr22 | 1,000,000 | 673 | 642 | 589 |
| Skok.NSD2 | chr22 | 2,500,000 | 141 | 139 | 136 |
| Skok.NSD2 | chrX | 10,000 | 4,310 | 3 | 0 |
| Skok.NSD2 | chrX | 25,000 | 34,214 | 7,574 | 987 |
| Skok.NSD2 | chrX | 50,000 | 42,908 | 13,666 | 11,329 |
| Skok.NSD2 | chrX | 100,000 | 32,173 | 14,589 | 5,103 |
| Skok.NSD2 | chrX | 250,000 | 27,555 | 6,623 | 6,346 |
| Skok.NSD2 | chrX | 500,000 | 19,387 | 6,640 | 6,043 |

*Continued on next page*

Table S7 – *Continued from previous page*

| **Dataset identifier** | **Chr.** | **Resolution** | **Non-empty blocks** | **IDR < 0.05** | **IDR < 0.01** |
|---|---|---|---|---|---|
| Skok.NSD2 | chrX | 1,000,000 | 9,628 | 4,364 | 4,182 |
| Skok.NSD2 | chrX | 2,500,000 | 1,957 | 1,418 | 969 |

Table S7: IDR2D analysis of individual chromosomes in three pairs of HiC experiments. Columns are (1) *chromosome identifier*, (2) *block resolution in base pairs*, (3) *number of non-empty (20 or more reads) blocks*, (4) *number of reproducible blocks with IDR* $< 0.05$, and (5) *number of reproducible blocks with IDR* $< 0.01$.
